# A Recombinant Protein Biomarker DDA Library Increases DIA Coverage of Low Abundance Plasma Proteins

**DOI:** 10.1101/2020.11.11.377309

**Authors:** Seong Beom Ahn, Karthik S. Kamath, Abidali Mohamedali, Zainab Noor, Jemma X. Wu, Dana Pascovici, Subash Adhikari, Harish R. Cheruku, Gilles J. Guillemin, Matthew J. McKay, Edouard C. Nice, Mark S. Baker

**Author notes:** Corresponding co-authors: Dr Seong Beom Ahn, Level 1, 75 Talavera Road, Macquarie University, 2109, Australia; Professor Mark S. Baker, Level 1, 75 Talavera Road, Macquarie University, 2109, Australia, Skype: msbaker377.

## Abstract

Credible detection and quantification of low abundance proteins from human blood plasma is a major challenge in precision medicine biomarker discovery when using mass spectrometry (MS). Here, we employed a mixture of recombinant proteins in DDA libraries to subsequently detect cancer-associated low abundance plasma proteins using SWATH/DIA. The exemplar DDA recombinant protein spectral library (rPSL) was derived from tryptic digestion of 36 human recombinant proteins that had been previously implicated as possible cancer biomarkers in both our own and other studies. The rPSL was then used to identify proteins from non-depleted colorectal cancer (CRC) plasmas by SWATH-MS. Most (32/36) of the proteins in the rPSL were reliably identified in plasma samples, including 8 proteins (BTC, CXCL10, IL1B, IL6, ITGB6, TGFα, TNF, TP53) not previously detected using high-stringency MS in human plasmas according to PeptideAtlas. The rPSL SWATH-MS protocol was compared to DDA-MS using MARS-depleted and post-digestion peptide fractionated plasmas (here referred to as a human plasma DDA library). Of the 32 proteins identified using rPSL SWATH, only 12 were identified using DDA-MS. The 20 additional proteins <u>exclusively</u> identified by using the rPSL approach with SWATH were mostly lower abundance (i.e., <10ng/ml) plasma proteins. To mitigate FDR concerns, and replicating a more typical approach, the DDA rPSL was also merged into a human plasma DDA library. When SWATH identification was repeated using this merged library, the majority (33/36) of low abundance plasma proteins from the rPSL could still be identified using high-stringency HPP Guidelines v3.0 protein inference criteria.

## Introduction

Biological fluids like plasma, serum, saliva and urine are commonly used for clinical diagnostic applications. Although plasma is highly heterogenous across a wide range of protein concentrations compared to many other biospecimens, it is a particularly attractive source for identification of disease-related protein biosignatures (1). Plasma collection is minimally invasive and hence readily available. It perfuses all tissues and has a relatively constant volume (~5 litres in an adult) and therefore has been suggested to represent patho-physiochemical snapshot of an individual at any given time.

However, the analysis of plasma proteins by tandem mass spectrometry (MS) is analytically challenging because of the presence of many “housekeeping” liver-derived proteins covering a large dynamic protein concentration range (>12 orders). For example, human serum albumin has a typical concentration of ~50mg/ml, whereas the cytokines IL-6 or IL-8 are present at the low pg/ml range (2). The masking of low abundance proteins (LAPs) by these highly abundant proteins (HAPs) makes it difficult to detect, identify and quantify many disease-specific protein biomarkers (3).

One approach to broaden the “reach” of plasma proteomics whilst mitigating masking effects involves removal of HAPs using immunodepletion, since the major 14 or 20 HAPs represent ~90 and ~97% of the total plasma proteome content, respectively (4). Another approach is to use extensive multidimensional peptide fractionation after plasma proteolytic digestion (5, 6) that facilitates the analysis of peptides from LAPs. Both methods are thought to allow the proteome to be explored in greater depth that better reflects pathophysiology, or reveal proteins that may be disease- or disease stage-specific (4, 7). Moreover, depleted and/or fractionated plasma samples are amenable to analysis by antibody-based technologies (8, 9) and/or MS (10). However, the use of depletion and fractionation methods independently has typically been limited to small pilot discovery studies (3) and is not yet amenable to automated high-throughput large clinical studies.

When shotgun proteomics is used to identify disease-related biomarkers (7), complex protein mixtures are routinely enzymatically digested or chemically cleaved, with subsequent peptides separated by HPLC followed by identification using tandem MS/MS. Utilising shotgun proteomics in combination with depletion or fractionation can allow the inference, identification and quantitation of ≥1000 plasma proteins from a single study (11, 12). Despite the high number of protein identifications using shotgun proteomics, disease-specific regulatory proteins expressed at extremely low levels frequently remain masked by HAPs.

Using a standard analytical pipeline, HUPO’s Biology/Disease-Human Proteome Project HPP (B/D-HPP) plasma proteome team has analysed 178 individual experiments. They have recently reported a total of 3,509 plasma proteins identified with imposition of community-endorsed high-stringency protein inference parameters. This HPP team also obtained evidence for an additional 1,300 proteins, although this was at lower stringency, less credible evidence (13), illustrating the difficulties associated with deep plasma proteome analyses.

Aside from established shotgun approaches, emerging MS technologies may have the potential to overcome some known limitations in protein identification and quantification. One such technology is Sequential Window Acquisition of All THeoretical Mass Spectra (SWATH MS), a data-independent acquisition (DIA) method that allows deep proteome coverage with the promise of comprehensive, accurate and reproducible quantitation (14). In SWATH-MS, all ionized peptides in the sample that fall within a specified mass range are fragmented in a systematic and unbiased fashion using precursor isolation windows (14). The resultant dataset constitutes a complete record of all individual peptides represented in a convoluted, but highly structured manner.

However, to achieve accurate quantification using SWATH-MS, it is crucial to have prior knowledge of the PSM (Peptide Spectrum Matches) and chromatographic behaviour of all peptides of interest. These PSMs can be used to extract peptide-specific information from observed MS spectral data using PQPs (Peptide Query Parameters) (14). This information includes peptide sequence, *m/z* values of the dominant precursor ion of the peptide, charge state, four to six most intense fragment ion *m/z* values of peptide(s) under fragmentation conditions, information about anticipated fragmentation and expected LC retention times.

These PQPs are commonly obtained from spectral data acquired from DDA (data-dependent acquisition) runs performed prior to a DIA experiment (14). The datasets employed are commonly referred to as “peptide spectral libraries”. Generating peptide spectral libraries using DDA is time-consuming, complicated and currently a major limitation of DIA MS, including SWATH (15, 16). Spectral libraries are usually equally, if not more, complex than the sample being analysed (although not a pre-requirement) and the quality of the data depends on factors such as isolation window widths, fragment ion resolutions, dwell times and cycle times (17). It is crucial to generate spectral libraries using an identical instrument type with identical settings when performing both DDA and DIA parts of all related experiments (16). Additionally, to minimise variation, a number of informatics tools have been developed and implemented (18). DIA specific software (e.g., PeakView^®^, Skyline) is then frequently used to identify unique peak groups associated with previously-generated DDA spectral libraries that can then be used to accurately identify and quantify target peptides observed in the DIA part of an experiment (16).

Given that SWATH/DIA is often reliant on previously generated DDA peptide spectral libraries (as opposed to library-free approaches), many studies have attempted to increase the depth of DDA libraries. Some have used a range of protein/peptide fractionation strategies (e.g., HAP depletion and/or combinations of peptide fractionation methods) prior to DDA experiments (4, 19). Other researchers have generated larger libraries by combining 2 or more existing libraries (20). Equally, software tools have been developed that facilitate library concatenation (15, 21). Clearly, generation of comprehensive, high-quality peptide spectral libraries for high-quality SWATH/DIA analysis is crucial, as these represent the biological/proteome space of biospecimens being interrogated.

Unfortunately, because of a requirement for libraries to be based on prior DDA experiments, the majority of SWATH/DIA plasma studies are unable to identify LAPs and, to our knowledge, no previous SWATH/DIA studies have been able to identify novel previously unidentified human plasma proteins. Rather, most choose to focus on protein quantitation of pre-identified library proteins. Although new tools like library-free DIA approaches (22) continue to be developed, library-based approaches remain widely used in SWATH/DIA applications. This study will focus on improving plasma protein depth and coverage outcomes using library-based approaches.

Here, we employ a novel approach to determine if it is possible to identify lower abundance cancer-associated plasma proteins using SWATH/DIA. A DDA spectral peptide library derived from tryptic peptides from a carefully-chosen set of 36 human recombinant proteins was constructed (Table 1). Each of the proteins selected had been previously implicated in human cancer/s and could be expected to be present (albeit possibly at low concentration) in human plasma. The previously reported plasma concentrations of those selected 36 proteins are summarised in Table 1. This set was based upon cancer biomarker protein candidates previously identified in the literature including BTC, C1QC, CEACAM5, CPQ, CXCL8, CXCL10, CXCL12, EGF, EGFR, IL1B, IL6, ITGAV, ITGB1, KLK3, MIA, MMP2/3/9, MUC1, PDGFB, PFN1, PLAU, PTEN, S100A8/9, TGFA, TIMP1/2, TNF, TNFRSF1A, TP53. In addition, our prior colorectal cancer studies implicated ADAMDEC1, CST3, ITGB6, PLAUR as potential early stage biomarkers and these have been combined with the literature biomarker candidates listed above to create the rPSL (4, 23–26). Importantly, 9 of the recombinant proteins used to construct this rPSL had <u>not</u> been previously identified in human plasma by MS (Table 1). The constructed recombinant protein spectral library (rPSL) was coupled to a SWATH-MS analysis (rPSL SWATH) workflow to determine if it was possible to observe tryptic peptide spectra for any of these 36 cancer-associated proteins in non-depleted, staged colorectal cancer (CRC) patient plasmas. The plasmas used in this study are identical to those employed by us previously (4, 26). Details of the experimental design are provided in Figure 1.

**Table 1.** Cancer-associated proteins (36 proteins) used to generate the rPSL DDA spectral library.

| Gene Name | Protein name | Human plasma concentration<sup>#</sup><br>(pg/ml – µg/ml) | Found in plasma by MS<sup>†</sup> |  | Cancers |  |
| --- | --- | --- | --- | --- | --- | --- |
|  |  |  | Y / N | Refs | Cancers | Refs |
| ADAMDEC1 | A disintegrin and metalloproteinase domain-like protein decysin-1 | 76 ng/ml | Y | (11, 29) | Colorectal | (4) |
| BTC | Probetacellulin | 4 ng/ml | N* | (30) | Ovarian | (31) |
| C1QC | Complement C1q subcomponent subunit C | 912 ng/ml | Y | (11, 29, 32, 33) | Prostate | (34) |
| CEACAM5 | Carcinoembryonic antigen-related cell adhesion molecule 5 | 1 ng/ml | Y | (11) | Colorectal, Lung, Gastric | (35-37) |
| CPQ | Carboxypeptidase Q | 29 µg/ml | Y | (11, 38, 39) | Liver | (40) |
| CST3 | Cystatin-C | 5 µg/ml | Y | (11, 41, 42) | Head & Neck | (43) |
| CXCL8 (IL8) | C-X-C motif chemokine 8 (Interleukin-8) | 14 pg/ml | N** | (44) | Colorectal, Brain, Breast | (45-47) |
| CXCL10 (IP10) | C-X-C motif chemokine 10 (Interferon gamma-induced protein 10) | 724 pg/ml | N** | (44) | Breast | (48) |
| CXCL12 (SDF-1α) | C-X-C motif chemokine 12 (stromal cell-derived factor 1) | 1 ng/ml | Y | (38, 49) | Head & neck, Oesophageal | (50, 51) |
| EGF | Pro-epidermal growth factor | 2 ng/ml | Y | (11, 39) | Lung | (52) |
| EGFR | Epidermal growth factor receptor | 12 ng/ml | Y | (11, 53) | Breast | (54) |
| IL1B | Interleukin-1 beta | 1 ng/ml | N** | (44) | Oral | (55) |
| IL6 | Interleukin-6 | 1 ng/ml | N** | (44) | Colorectal, Prostate | (56, 57) |
| ITGAV | Integrin alpha V | 2 ng/ml | Y | (38, 49) | Ovarian | (58) |
| ITGB1 | Integrin beta 1 | 2 µg/ml | Y | (11, 38) | Breast | (59) |
| ITGB6 | Integrin beta 6 | 2 ng/ml | N* | (30) | Colorectal | (58, 60) |
| KLK3 | Plasma kallikrein | 26 ng/ml | Y | (33, 38) | Prostate | (61) |
| MIA | Melanoma-derived growth regulatory protein | 9 ng/ml | Y | (49) | Melanoma | (62) |
| MMP2 | Matrix metalloproteinase 2 | 812 ng/ml | Y | (49, 53, 63) | Colorectal, | (64, |
|  |  |  |  |  | Breast | (65) |
| MMP3 | Matrix metalloproteinase-3 /Stromelysin-1 | 79 ng/ml | Y | (11, 38) | Ovarian | (66) |
| MMP9 | Matrix metalloproteinase 9 | 190 ng/ml | Y | (11, 38, 53) | Colorectal, Breast | (64, 65) |
| MUC1 | Mucin-1 | - | Y | (13) | Breast | (67) |
| PDGFB | Platelet-derived growth factor subunit B | 323 pg/ml | Y | (11, 38) | Colorectal | (68) |
| PFN1 | Profilin-1 | 129 ng/ml | Y | (38, 39) | Lung | (69) |
| PLAU | Urokinase-type plasminogen activator | 794 pg/ml | Y | (11) | Prostate | (57) |
| PLAUR | Urokinase plasminogen activator surface receptor | 2 ng/ml | Y | (11) | Colorectal, Lung, Prostate | (70-72) |
| PLG | Plasminogen | 302 µg/ml | Y | (11, 33, 38, 39) | Ovarian | (73) |
| PTEN | Phosphatase tensin homolog | 295 pg/ml | Y | (11, 38, 39) | Prostate | (74) |
| S100A8 | Protein S100-A8 | 269 ng/ml | Y | (38, 39) | Bladder | (75) |
| S100A9 | Protein S100-A9 | 2 µg/ml | Y | (38, 39) | Bladder | (75) |
| TGFA | Pro-transforming growth factor α | 15 pg/ml | N* | (30) | Gastric | (37) |
| TIMP1 | Metalloproteinase inhibitor 1 | 269 ng/ml | Y | (32, 38, 41) | Colorectal, Breast | (76, 77) |
| TIMP2 | Metalloproteinase inhibitor 2 | 151 ng/ml | Y | (11, 29, 49) | Colorectal | (78) |
| TNF | Tumour necrosis factor | 2 ng/ml | N*** | (79) | Breast | (80) |
| TNFRSF1A | Tumour necrosis factor receptor superfamily member 1A | 5 ng/ml | Y | (11) | Prostate | (81) |
| TP53 | Cellular tumour antigen p53 | 398 pg/ml | N** | (44) | Bladder | (82) |

**Figure 1.**
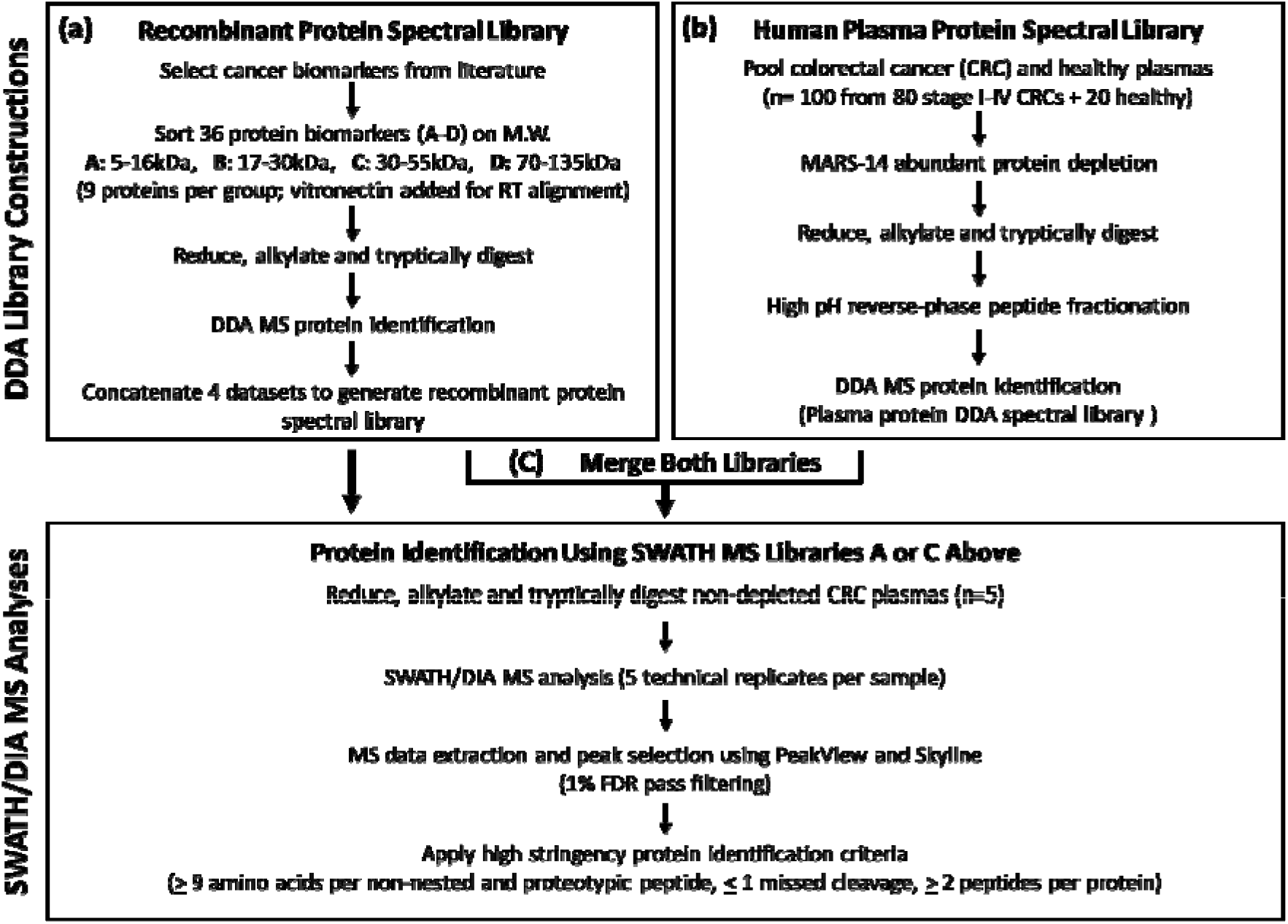
Experimental workflow. **(a) Construction of recombinant protein spectral library**. A total of 36 cancer-associated biomarkers were selected from the literature and our own studies (Table 1) and sorted into 4 groups (A - D) based on their molecular weight (M.W.). Each group of 9 proteins was spiked with vitronectin for retention time (RT) alignment. Proteins were reduced, alkylated and digested with trypsin. DDA was used for protein identification using a SCIEX TripleTOF 6600. Datasets were concatenated to generate a recombinant protein spectral library. **(b) Construction of human plasma protein spectral library.** CRC plasma (80 from stage I-IV CRCs) and 20 healthy plasma samples were pooled and depleted the top 14 high abundance proteins with an Agilent MARS-14 depletion column. The depleted samples were digested with trypsin followed by peptide fractionation using high pH reverse-phased HPLC. A DDA method was employed as in (a) to construct the plasma protein DDA spectral library. **(c) SWATH/DIA protein identification using rPSL or merged libraries.** The constructed rPSL or merged libraries (rPSL + plasma protein spectral library) were combined with SWATH/DIA MS analysis to determine whether it was possible to detect tryptic peptide spectra of the 36 cancer-associated proteins in non-depleted human plasma samples obtained from CRC patients (n=5). Following sample preparation (reduction, alkylation and tryptic digestion), SWATH/DIA MS analysis was performed for peptide/protein identification. PeakView and Skyline were employed for MS data extraction and peak selection with 1% FDR filtering. Identified proteins were further filtered using high stringency protein identification criteria (HPP guideline v3.0).

To demonstrate the ability of rPSL SWATH to detect low abundance cancer-associated proteins from non-depleted plasma, we compared protein identifications from rPSL SWATH to previous plasma protein identification methods (i.e., DDA shotgun proteomics using MARS-14-depleted or peptide fractionated tryptically-digested CRC plasma proteins, referred to as the “plasma protein DDA spectral library”).

To mitigate FDR concerns due to the use of a small 36 recombinant protein rPSL, the rPSL was merged with an additional human plasma protein DDA spectra library (containing 742 plasma proteins) and SWATH-MS protein identification performed using HPP Guidelines v3.0 (i.e., high stringency identification criteria of ≥ 2 peptides non-nested, uniquely-mapping peptides of ≥ 9 amino acids length and limited to ≤ 1 missed cleavage) (27, 28).

Our approach indicates that rPSL SWATH allows identification of many very low abundance, previously undetectable, plasma proteins in the pg - ng/ml range. rPSL SWATH therefore appears to have the potential to discover low abundance plasma proteins that potentially have utility as diagnostic, prognostic or theranostic indicators for cancer or detection and surveillance of other disease.

## Materials and Methods

### Ethics statement and plasma sample collection

This study was performed with approval #5201200702 from Macquarie University Human Research Ethics Committee. The EDTA-plasma cohort contained 100 individuals patients (80 clinically-staged CRC patients and 20 healthy controls) procured from the Victorian Cancer Biobank, Melbourne, Australia. Sample details and preparation methods have been described previously (4, 26).

### Recombinant proteins

Recombinant proteins ADEMDEC1 (TP721090), BTC (TP723036), C1QC (TP761200), CPQ (TP760108), CXCL10 (IP10, TP723726), CXCL8 (IL8, TP721122), MMP2 (TP723320), MUC1 (TP760771), PDGFB (TP723355), TGFA (TP723858), TIMP2 (TP723886) and TNFRSF1A (TP723870) were purchased from OriGene Technologies, CEACAM5 (4128-CM), CST3 (1196-PI), CXCL12 (350-NS), EGFR (1095-ER), IL1B (201-LB), IL6 (206-IL), KLK3 (1344-SE), MIA (9250-IA), MMP3 (513-MP), MMP9 (911-MP), PLAU (1310-SE), PLAUR (807-UK), PTEN (847-PN), S100A8/A9 (8226-S8), TIMP1 (970-TM),TNF (210-TA), TP53 (SP-454) and VN (2308-VN) from R&D Systems, PFN1 (NBP1-30215), PLG (H00005340-P01), ITGAV (H00003685-P01), ITGB1 (H00003688-P01) and ITGB6 (H00003694-P01) from Novus Biologicals and EGF (MBS650012) from My BioSource.

### Sample preparation

For rPSL construction, selected recombinant proteins were pooled (Table 1) into four groups based on similar molecular weight (MW) (namely, groups A: 5-16kDa, B: 17-30kDa, C: 30-55kDa and D: 70-135kDa). Samples were reduced with 5mM dithiothreitol (DTT) at 60°C for 30min followed by alkylation with 25mM iodoacetamide (IAA) at room temperature for 30min in the dark. Samples were then digested with sequencing grade porcine trypsin (Promega) at a protease to substrate ratio of 1:30 at 37°C for 16hr. Peptide mixtures were desalted and cleaned with C18 OMIX tips (Agilent) according to the manufacturer’s protocol, followed by drying using vacuum centrifugation.

For the human plasma library construction, all 100 plasma samples were pooled and immunodepleted using an Agilent MARS-14 high capacity affinity column with a Agilent 1260 HPLC system as described previously (4). The depleted plasma samples were then reduced, alkylated and digested as described. Digested peptides were fractionated (total 20 fractions) using a ZORBAX 300 Extend-C18 (2.1×150 mm, 3.5μm) column on a 1260 HPLC system (Agilent, Santa Clara, CA, USA) as described previously (4). Fractioned peptides were desalted and cleaned with C18 OMIX tips and dried by vacuum centrifugation.

### Spectral library generation (DDA)

The rPSL (groups A-D above) and the human plasma spectral library (using 20 fractionated peptides as described above) were generated independently. DDA protein identification was performed on a SCIEX TripleTOF 6600 (SCIEX, Framingham, MA) coupled to an Eksigent Ultra nanoLC system (Eksigent Technologies, Dublin, CA). Peptides were injected onto 200μm ID, 3.5-cm-length peptide-trap columns packed in house with a C18 support (2.7μm particle size, Halo C18) for pre-concentration and desalted at a flow rate of 5μL/min for 3min with 0.1% formic acid (v/v) and 2% acetonitrile (v/v). After desalting, the peptide trap was switched in-line with a cHiPLC C18 column (15 cm x 200μm, 3μm, ChromXP C18-CL, 120 Å, 25°C, SCIEX) and peptides were eluted using a linear 120min gradient from 5 % acetonitrile to 35 % mobile phase B (B: 99.9 % acetonitrile, 0.1 % formic acid) at 600 nL/min. In DDA mode, a TOFMS survey scan was acquired at m/z 350 – 1500 with 0.25 second accumulation time, with the 20 most intense precursor ions (2+-5+; counts >200) in the survey scan consecutively isolated for subsequent product ion scans. Dynamic exclusion was used with a window of 30secs. Product ion spectra were accumulated for 100msecs in the mass range m/z 100 – 1800 with rolling collision energy.

DDA data were analysed using ProteinPilot (V5.0, SCIEX) with the Paragon algorithm (84). The *Homo sapiens* protein sequence database with reviewed entries was obtained from SwissProt (42,388 entries including canonical proteins and isoforms, 2018 version). The search parameters were as follows: sample type: identification; cys alkylation: iodoacetamide; digestion: trypsin; instrument: TripleTOF 6600; ID focus: biological modifications; precursor peptide mass tolerance: ±50ppm. A reverse-decoy database search strategy was used with ProteinPilot, with the calculated protein at 1% FDR and a detected protein threshold [Unused ProtScore (Conf)] >:1.30 (95.0%). The data has been deposited to ProteomeXchange Consortium via the PRIDE partner repository (PXD022361).

### *In silico* peptide repertoire of recombinant proteins

Recombinant proteins were digested *in silico* to identify uniquely-mapping non-nested peptides of minimum length nine amino acids, similar to our previous *in silico* analysis (85). Peptide uniqueness (i.e., unitypic peptides) was confirmed using the neXtProt uniqueness checker (86).

### High stringency peptide/protein selection

HPP guidelines v3.0 were applied for the application of high stringency protein inferences (27). Maximum of 1 missed cleavage was allowed for each unitypic and non-nested peptide, with tryptic cleavage towards the C-terminal side of the R/K residues but not when immediately followed by a Proline residue. Greater than or equal to 2 peptides per protein was required for protein identification.

### Spectral library merging

The independently generated rPSL and human plasma spectral library were merged into one spectral library using previously published algorithm, SWATHXtend [25]. The rPSL was used as the seed library when merging. No modifications and miscleaved peptides were removed. Only peptides with confidence >0.99 were considered. The matching quality between the two libraries is excellent with the squared retention time correlation of 0.97, estimated retention time error of 1.74min and relative ion intensity correlation of 0.9. The merged spectra library was outputted in a tab-delimited PeakView compatible text format.

### DIA/SWATH-MS

A SCIEX TripleTOF 6600 coupled with Eksigent Ultra nanoLC system with identical mobile phase conditions to those described above was used for SWATH-MS. For SWATH, data was acquired using a shorter 60min LC gradient (5-35% mobile phase B) at 600nl/min. The eluent was subjected to positive ion nanoflow electrospray MS analysis. Initially, the precursor m/z frequencies from previously generated plasma proteome DDA data were used to determine the m/z window sizes. SWATH variable window acquisition with a set of 100 overlapping windows was constructed covering the mass range m/z 400 – 1000. In SWATH mode, TOF-MS survey scans were acquired (m/z 350-1800, 0.05 sec) then the 100 predefined m/z ranges were sequentially subjected to MS/MS analysis. Product ion spectra were accumulated for 30msecs in the mass range m/z 200-2000 with rolling collision energy optimized for lowed m/z in m/z window +10%.

### DIA/SWATH Peak Extraction

DIA spectral alignment and targeted data extraction were performed independently using two software packages; Skyline software (Version 4.1.0.18169) (87) and PeakView (SCIEX, USA). In both instances the same DDA based spectral library (~950 peptides and ~23,000 transitions) was utilized to execute the targeted data extractions. DIA chromatograms were extracted from SWATH-MS data files (*n*=5 per condition: replicates indicate the technical injection replicates).

### Peak extraction using PeakView

Ion library and SWATH data files were imported into PeakView 2.1 with SWATH quantitation plug-in (SCIEX). Retention times for all CRC sample SWATH data files were aligned using linear regression by selecting 5 endogenous peptides across the elution profile. The top 6 fragment ions for each peptide were extracted from the SWATH data using 75ppm target XIC width, peptide confidence threshold of ≥ 0.99, and a 10min retention time extraction window. After data processing, peptides with confidence >99% and FDR <1% (based on chromatographic feature after fragment extraction) were used for quantitation. Shared and modified peptides were excluded. The sum of MS2 ion peak areas of SWATH quantified peptides for individual proteins were exported to calculate the protein peak areas.

### Peak extraction using Skyline

To import SWATH data, isolation window schemes ranging from 399 m/z to 1000 m/z were extracted from data files. To perform retention time (RT) calibration and to design a RT predictor, an indexed retention time (iRT) calculator using 12 high intensity endogenous vitronectin and human serum albumin peptides with retention times spanning the whole gradient were used.

A list of target proteins was generated by importing the ion library which was filtered to remove duplicate peptides and peptides of <7 amino acids in length. For targeted peptides, the top-ranked six *y* and *b* transition ions with charges up to +3 were selected together with their corresponding decoy-transition groups, generated by shuffling sequences from the targeted peptides. Detailed peptide and transition settings are provided in supplementary table S1. Chromatograms were extracted directly from the raw data files within a window of 20min (± 10min) around the predicted times (with a resolving power of 30,000 for TripleTOF 6600). On inspection of the extracted peaks, it was noted that some of the peaks appeared slightly outside of the narrow 10min window. The window size was therefore increased to 20min to include these refined manually inspected peaks.

For SWATH-MS data, chromatographic peaks were integrated using mProphet (88) peak-scoring model, which is a semi-supervised learning algorithm, to identify correct peaks. Peak scoring models were refined using feature scores of peptides, estimating Q-values for each peak. A false discovery rate (FDR) of 1% (Q value cut-off 0.01) was applied and peaks were filtered by applying a dot product (dotP) cut-off of 0.6 which shows the correlation between observed (library) and measured (DIA) spectra.

Peptide and protein intensities were calculated by manually summing average peak areas of respective MS/MS transitions and peptides, respectively. Subsequently, filters to remove standard RT calibration peptides and peptides with ≥2 missed cleavages were applied for selecting unique peptide candidates.

### Data processing

SWATH extractions by Skyline and PeakView initially went through data cleaning and filtering before comparisons. All decoy peptides, RT-calibration peptides and peptides with more than two mis-cleavages were removed. All modified peptides were removed except carbamidomethylated cystine and oxidised methionine containing peptides. The data subsequently was further filtered using FDR criteria. Two different FDR criteria were used for the SWATH data extracted using the two different (rPSL and plasma) libraries. For the SWATH data extracted using the merged spectral library, the default PeakView FDR criterion was used, i.e., peptides with at least one sample satisfying FDR < 0.01 were kept (15, 21). For the SWATH data extracted using the rPSL, a more stringent FDR criterion (>3 replicates within a group having FDR < 0.01) was applied due to the potential for less effective FDR calculation resulting from the use of a small spectral library.

## Results

### Construction of a cancer-associated recombinant protein spectral library (rPSL)

Data processing and peak feature extraction for identification of proteins through SWATH-MS analysis is dependent on the quality of previously generated DDA spectral libraries (15, 16). The quality and coverage of spectral libraries has been found to be directly associated with the efficacy and scope of finding potential candidates from any SWATH-MS analyses (89).

In this study, a rPSL was employed to assist in identifying lower abundance cancer-associated plasma proteins by SWATH-MS (Fig 1). In the initial experiment, a high-quality rPSL with broad coverage for all of the 36 full-length recombinant proteins, each of which has previously been reported as a plasma cancer biomarker, was generated (Table 1). The rPSL proteins were selected strictly based on literature experimental evidence (e.g., MS, ELISA, Western Blotting). Analysis using PeptideAtlas confirmed 9 of these proteins (BTC, CXCL8, CXCL10, IL1B, IL6, ITGB6, TGFα, TNF, TP53) had not previously been detected in human plasma by MS. The plasma concentration of these proteins has been reported to be <10ng/ml (Table 1), reflecting the inherent challenge of low abundance plasma protein identification by MS in the presence of HAPs.

rPSL generation was performed using DDA. As expected, all 36 recombinant proteins were detected, generally through a high number of peptides identified at high confidence (≥ 99%). The highest peptide number detected in the rPSL was for integrin ITGAV (102 peptides reflecting 64% ITGAV coverage) while the lowest was CST3 (10 peptides and 36% coverage). Twenty-eight (28) proteins out of the 36 chosen had greater than 40% protein coverage. EGF had the highest coverage (100%) with mucin MUC1 having the lowest coverage (12%) (Table 2 and Supp. Table S2).

**Table 2.**
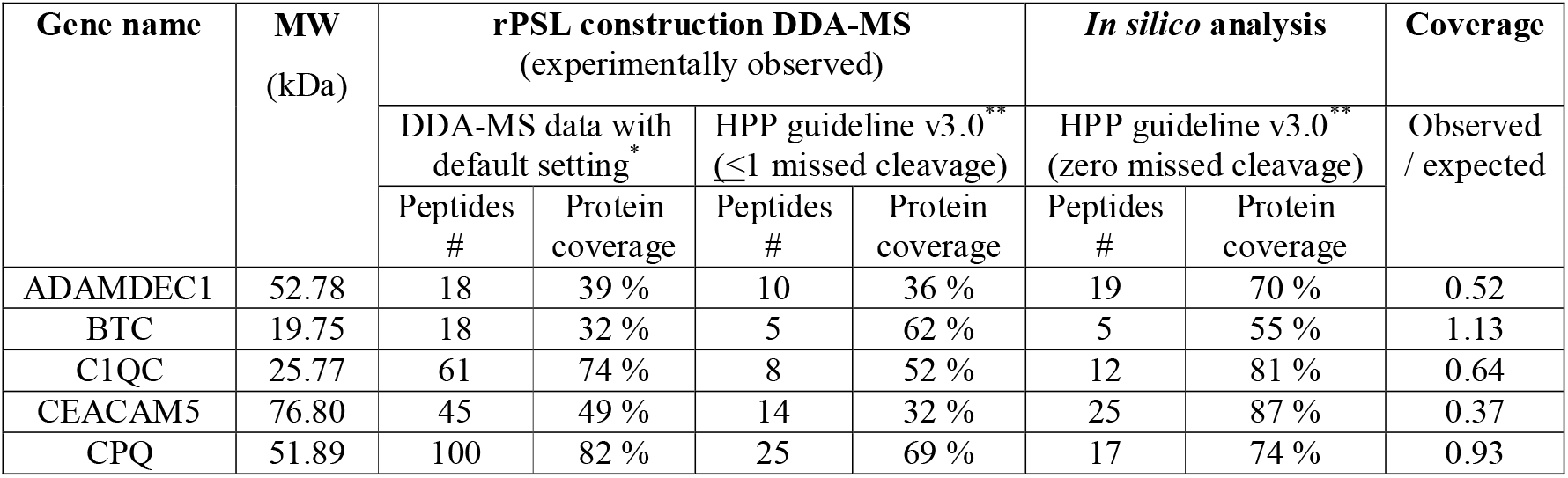
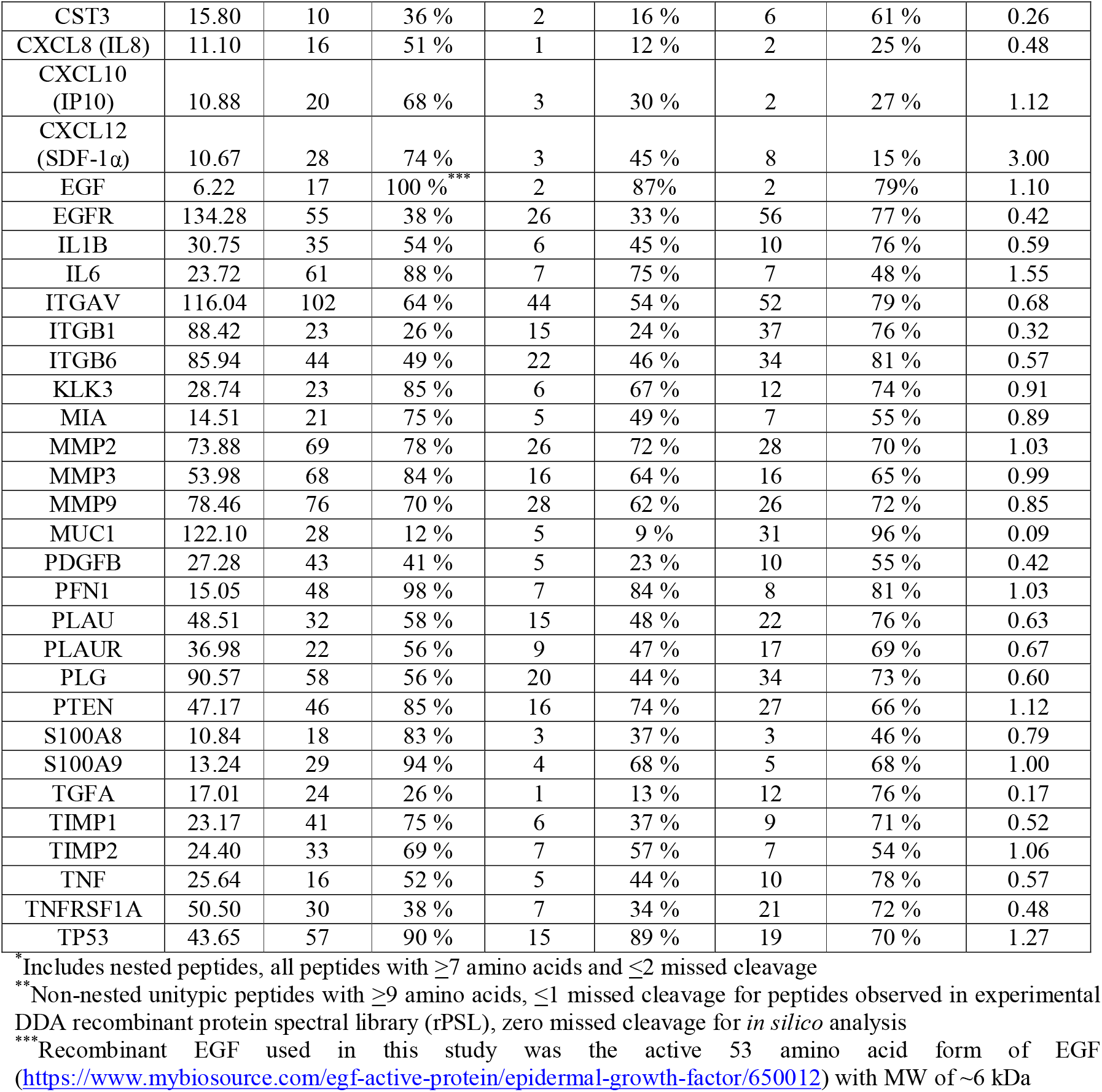
Peptides detected and protein coverage for each rPSL recombinant protein used to construct the experimental DDA spectral library and theoretical *in silico* analysis. Supplementary Table S2 contains details including sequences for all peptides.

To facilitate development of subsequent targeted assays (SRM, MRM or PRM) for future biomarker studies, we undertook a comparative analysis on experimentally observed uniquely-mapping (unitypic) peptides against a unitypic peptide repertoire derived *in silico* from the 36 recombinant proteins sequences. Identified peptides were filtered to select only unitypic peptides satisfying current HPP Guidelines v3.0 (see Table 2), including ≤1 missed cleavage. Concurrently, an *in silico* tryptic digestion analysis was performed for each of the 36 proteins with the same stringency guidelines applied except a zero missed cleavages rule was imposed. Proteome coverage corresponding to these *in silico* peptides for each protein was calculated to derive the maximum theoretical sequence coverage utilising tryptic peptides.

As expected, the number of identified peptides observed and protein coverage of the recombinant proteins both decreased when HPP high-stringency protein inference metrics were applied, primarily due to the removal of nested, non-unitypic and/or >1 missed cleavage peptides (Table 2). When compared to *in silico* data, some proteins (i.e., COQ, MMP2/3, KLK3, ITGB6 and S100A9) had a high number of peptide identifications as well as high coverage (see Table 2; coverage observed/expected). This indicates that most unitypic peptides satisfying high-stringency HPP MS Guidelines v3.0 requirements could be observed in the rPSL.

For some proteins, a higher number of peptides and coverage were observed experimentally at HPP stringency compared to *in silico* expectations. This is likely because of the zero missed cleaved rule applied for *in silico* analysis, whereas ≤1 missed cleavage was applied for the experimentally observed peptides (i.e., the DDA generated rPSL).

Furthermore, high-stringency *in silico* prediction suggested that only 2 potentially detectable peptides may be available for both CXCL10 (IP-10) and CXCL8 (IL-8). The plasma concentrations of these proteins are low (both in pg/ml range) and neither have been previously identified from plasma by MS (Table 1). Our *in silico* MS analysis provides a possible explanation why some plasma proteins have not yet been identified in human plasma. However, our novel rPSL approach detects peptides at the high stringency level (3 for CXCL10 and 1 for CXCL8) that has allowed identification of these very low abundance plasma proteins from non-depleted CRC plasma samples (see section below).

Table 2 suggests that our rPSL approach produces a more comprehensive library that significantly facilitates the challenge of detecting peptides from low abundance proteins in complex samples by SWATH/DIA.

### CRC plasma protein biomarker identification using the rPSL SWATH approach

In order to identify how many lower abundance cancer-associated plasma rPSL proteins were present in CRC plasmas, non-depleted plasmas from pooled CRC patients (n=5; see Fig 1) were examined by SWATH. In detail, instead of using SWATH in the traditional manner (protein quantification), we repurposed it to detect peptides that would ordinarily not be captured in standard plasma DDA experiments. This novel approach allowed low abundance proteins to be detected in a single experiment, importantly without depletion or extensive peptide fractionation. Initially, a SWATH-MS dataset was generated from each CRC plasma sample undertaken through 5 technical replicates. SWATH-MS datasets were analysed with two independent DIA analysis software tools, namely PeakView (PV) and Skyline (SL).

Of the 36 rPSL biomarkers, we reliably identified 32 proteins using PV and 23 proteins using SL in CRC plasmas. In most cases, higher peptide counts were observed from PV compared to SL (Supp. Table S3). Although proteins identified from PV covered all proteins from SL, we noted that half of the SL identified peptides were not common to PV. This was expected as each software package relies on different algorithms for detection and quantification of unique peptides and proteins. Similar discordances have been observed in a previous study (90). The purpose of using two different software packages was primarily to test whether the rPSL SWATH approach allowed detection of low-abundance cancer-associated proteins through either of the PV and SL platforms. Our study has not focused on a comprehensive comparison of the PV and SL software tools, and readers should refer to a previous article (90) for such a comprehensive multicentre benchmarking study involving label-free proteome quantification. To visualize the detectable threshold of low abundance proteins using the rPSL SWATH approach, we superimposed the 32 proteins identified onto a human plasma protein concentration curve (Fig 2). This concentration curve has been created in-house using reported plasma concentrations obtained from the Plasma Proteome Database (83), PeptideAtlas and comprehensive literature searches.

**Figure 2.**
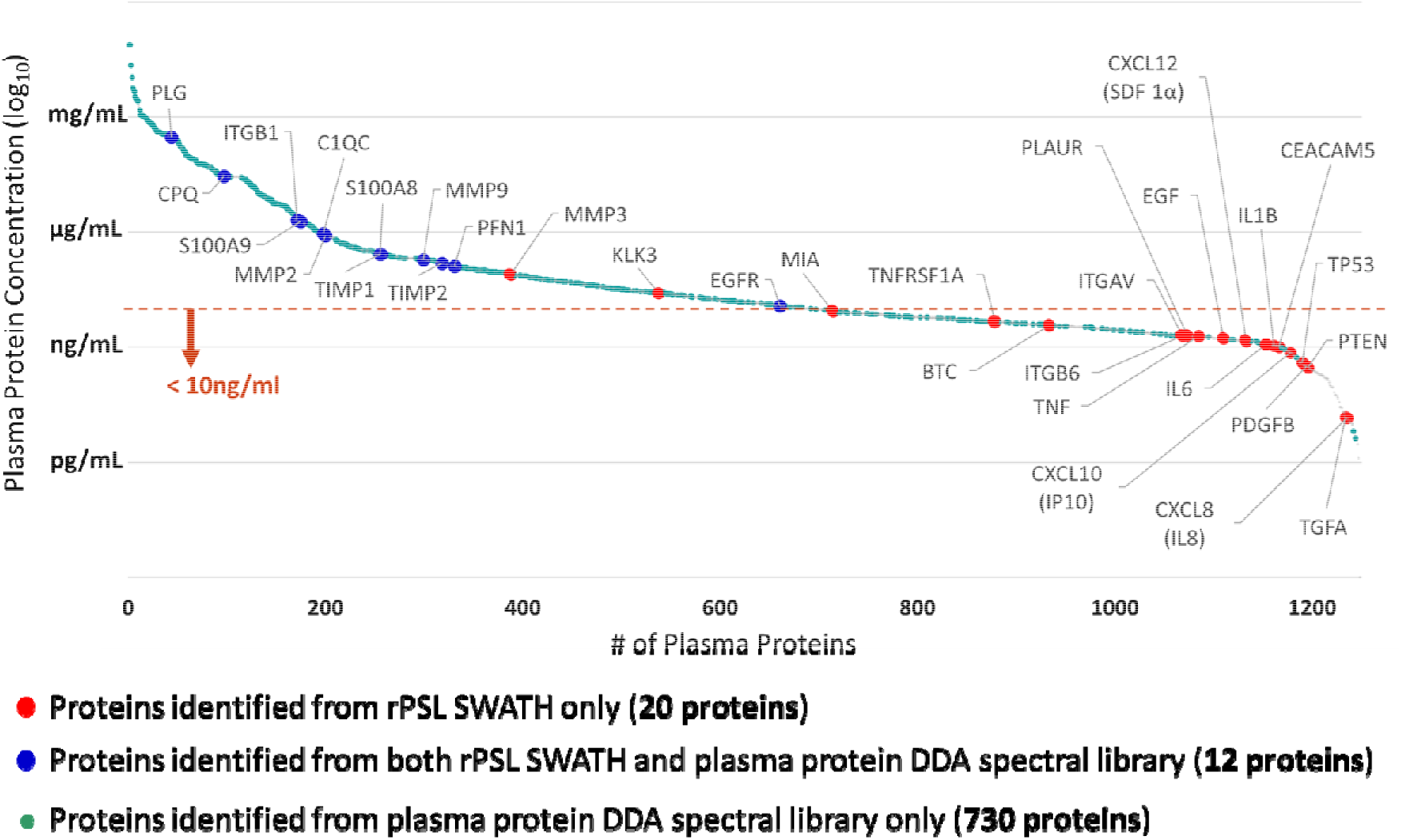
A superimposition of plasma protein concentrations of proteins identified from the recombinant protein spectral library coupled with SWATH-MS analysis (rPSL SWATH) and the plasma protein DDA spectral library. Supplementary Table S4 contains a list of all identified plasma proteins and peptide sequences using the DDA spectral library.

One interesting observation from this data is that rPSL SWATH allows identification of candidate biomarkers that are widely spread across the plasma protein concentration range (i.e., high through to low abundance proteins, Fig 2). Most importantly, rPSL SWATH allowed detection of plasma proteins BTC, CXCL8, CXCL10, IL1B, IL6, ITGB6, TGFα, TNF and TP53 that to the best of our knowledge have not been previously detected in human plasma using any MS technology. Furthermore, the data strongly support the hypothesis that low abundance cancer-associated plasma proteins present in the pg/ml range (CEACAM5, CXCL8, CXCL10, PDGFB, PTEN, TGFA, TP53) can be detected using non-immunodepleted patient plasma samples.

To demonstrate the ability of rPSL SWATH to detect low abundance cancer-associated proteins from non-depleted CRC plasma, we compared protein identifications from the rPSL SWATH to our routine plasma protein identification method (i.e., DDA shotgun proteomics on HAP-depleted or peptide fractionated plasma, referred to as “plasma protein DDA spectral library”, see Fig. 1).

To maximise plasma protein identifications from DDA shotgun runs, we combined healthy/CRC plasma samples (n=100) to ensure proteins present in both healthy and disease conditions were present. We subsequently removed the top 14 HAPs using Agilent’s MARS-14 system. Following tryptic digestion of MARS-14-depleted plasmas, reversed-phased hydrophobic interaction high pH fractionation (4) was employed to separate peptides into 20 fractions. For protein identification, we used an identical MS instrument and settings as for the rPSL SWATH. We were able to identify a total of 742 plasma proteins from pooled healthy or CRC plasmas (Fig 2 and Supp. Table S4 for a full list of identified proteins and peptide sequences). Compared to our previous study (4), we identified an additional 229 plasma proteins using the more advanced SCIEX TripleTOF 6600 MS instrument (previously 513 proteins identified from identical samples using a Sciex TripleTOF 5600) (4).

Of the 32 cancer-associated plasma proteins identified using rPSL SWATH, 12 proteins could also be identified in this DDA plasma library (Fig 2). Interestingly, the reported concentrations of these 12 proteins were generally higher abundance proteins (Fig 2, blue dots on plasma protein concentration curve). The remaining 20 proteins could exclusively only be identified by our novel rPSL SWATH approach, and these were generally lower abundance proteins (<10 ng/ml; with exception of MMP3 and KLK3; Fig 2, red dots).

### CRC plasma protein biomarker identification using a merged rPSL <u>and</u> human plasma protein DDA spectral library

Although SWATH analysis is one of the most advanced MS technologies, there are still some concerns regarding inconsistency of data analyses from a single library compared to merged libraries (91). One of the inevitable issues is FDR correction when merging different sized DDA libraries for the SWATH analysis. To mitigate FDR concerns resulting from use of a small single library (like rPSL), we merged the 36 protein rPSL library with a more standard plasma protein DDA library (containing data for 742 plasma proteins). SWATH analysis was then performed with identical experimental settings as used above (Fig 1). The main reason for using a merged library was to determine if SWATH continued to allow identification of candidate low abundance biomarker proteins in the presence of a much more complex library background.

We compared lists of identified proteins and peptides (i.e., quantifiable proteins and peptides) derived from rPSL SWATH and SWATH MS analysis using the merged library. RT alignments between libraries, as well as the CRC plasma SWATH MS dataset, were pegged against peptides derived from the abundant and ubiquitous proteins vitronectin and albumin. SWATH MS on the merged library identified a total of 519 proteins supported by 3,187 quantifiable peptides (Fig 3, Supplementary Table S5). All the previously observable 32 proteins biomarkers identified using the rPSL SWATH method could also be identified using a merged rPSL+ depleted plasma DDA library. In many cases, a greater number of peptides were identified using the merged library method compared with the rPSL SWATH solely (Fig 3a). Similar numbers of peptides were identified for 3 proteins (BTC, CXCL8, TIMP2), and for one protein (PTEN) a higher number of peptides (4 peptides) were identified from rPSL SWATH compared to the merged library SWATH (3 peptides) (Fig 3a). When comparing the total number of identified peptides, 47 peptides were common across both methods, and 87 peptides were unique to rPSL (Fig 3b). A list of identified peptides from rPSL SWATH and merged library SWATH is presented in Supplementary Table S6.

**Figure 3.**
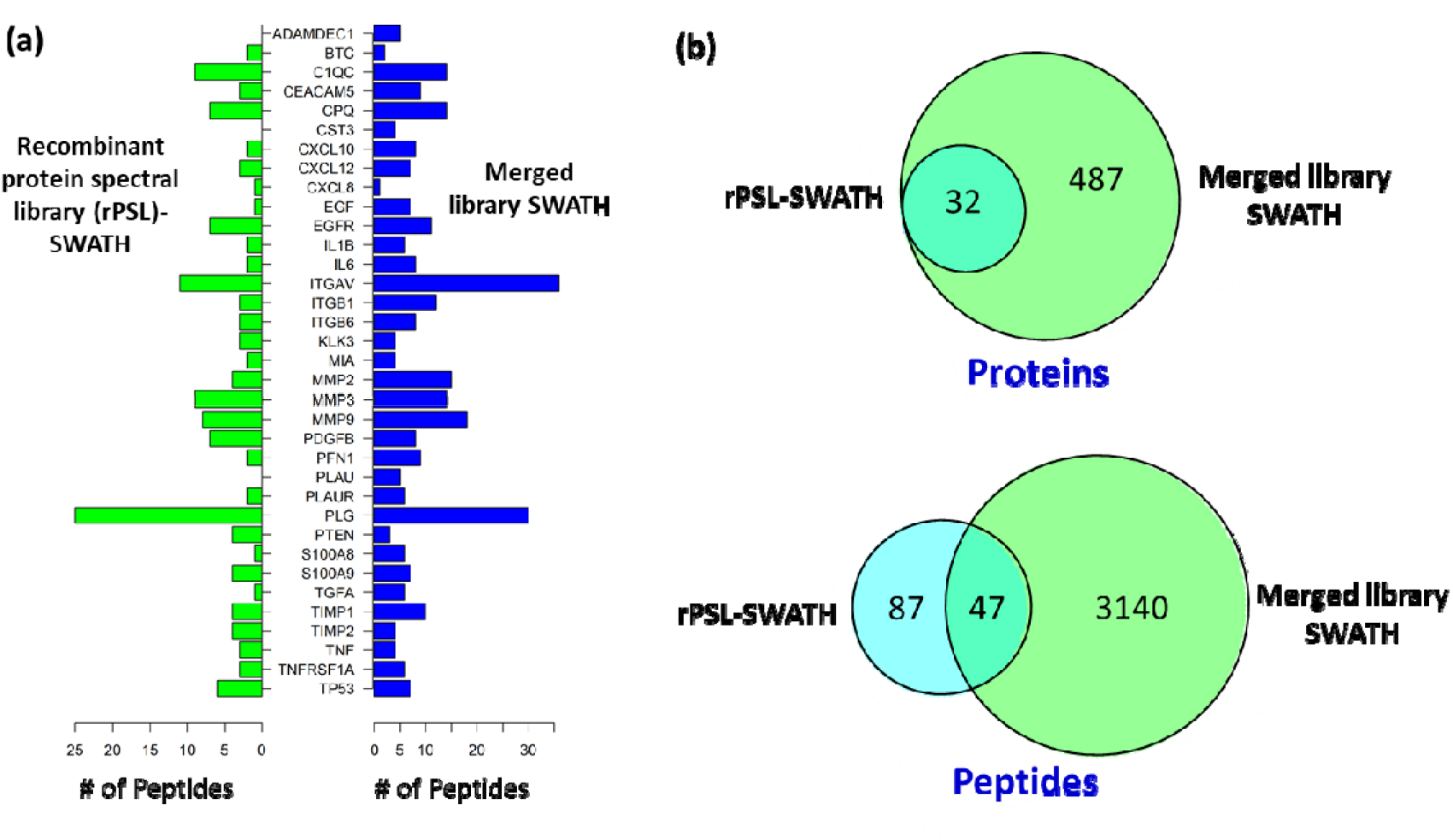
A comparison of identified plasma proteins and peptides between recombinant protein spectral library coupled with SWATH-MS analysis (rPSL SWATH) and SWATH-MS analysis on the merged library. **(a)** Number of detected peptides from rPSL SWATH and SWATH-MS on merged library demonstrating a similar trend but showing 3 additional proteins identified in the merged approach. **(b)** Venn diagrams comparing the number of common and unshared identified proteins and peptides between rPSL SWATH and SWATH-MS on merged library. A list of identified peptides from rPSL SWATH and merged library SWATH is presented in Supplementary Table S6.

Although the overall trend of numbers of peptides identified was relatively consistent across both approaches, we were able to credibly identify 3 additional proteins (ADAMDEC1, CST3 and PLAU) using the merged library. This increases the coverage using the rPSL from 32 (rPSL) to 35 (rPSL + depleted plasma DDA library) out of the original 36 recombinant proteins employed in this study. Interestingly, the identified peptides from the extra three proteins originated from the rPSL part of the merged library, not the depleted plasma DDA spectral library (Supplementary Table S7). We assume this is likely due to computational issues around FDR (92) and this possibility is discussed further below.

To increase peptide identification confidence, we finally applied high-stringency protein inference criteria (HPP Guidelines v3.0) (27) to all detected peptides using the merged rPSL + depleted plasma DDA library (See Fig 1 for details). Not surprisingly, the number of identified peptides significantly reduced for each protein after high-stringency filtering (Table 3). Interestingly, CXCL8 and TIMP2 were disqualified as they only had one peptide identified. Supplementary Table S8 contains data on the proteins and peptide sequences before and after applying high stringency protein identification criteria.

**Table 3:** Detected peptides and proteins from non-depleted CRC plasma by SWATH-MS analysis using merged library (rPSL + plasma protein spectral library). The table only contains unitypic peptides that satisfied the high-stringency protein inference/identification criteria, HPP Guideline v3.0. See supplementary Table S8 for more details.

| Gene names | Peptides #<br>(HPP Guidelines v3.0) * | Gene names | Peptides #<br>(HPP Guidelines v3.0) * |
| --- | --- | --- | --- |
| ITGAV | 22 | IL1B | 3 |
| PLG | 13 | IL6 | 3 |
| MMP9 | 11 | MIA | 3 |
| MMP2 | 10 | PDGFB | 3 |
| MMP3 | 9 | TNFRSF1A | 3 |
| ITGB1 | 9 | CST3 | 2 |
| TIMP1 | 7 | CXCL10 (IP10) | 2 |
| PFN1 | 7 | CXCL12 (SDF-1α) | 2 |
| CPQ | 7 | EGF | 2 |
| CEACAM5 | 6 | BTC | 2 |
| ITGB6 | 6 | PTEN | 2 |
| TP53 | 6 | S100A8 | 2 |
| C1QC | 5 | S100A9 | 2 |
| EGFR | 5 | TGFA | 2 |
| PLAU | 5 | TNF | 2 |
| ADAMDEC1 | 4 | CXCL8 (IL8) ** | 1 |
| KLK3 | 4 | TIMP2 ** | 1 |
| PLAUR | 4 |  |  |

\* non-nested untypic peptides with $\geq 9$ amino acids, $\leq 1$ missed cleavage, $\geq 2$ peptides per protein
\*\* proteins that were not qualified with high stringency protein identification criteria

## Discussion

Identification of early-stage, low abundance human plasma cancer biomarkers is an objective of many biomarker studies. Recently, advanced technologies like liquid biopsies (i.e., detecting ctDNA) (93) and SWATH/DIA-MS (i.e., identifying and quantitating cancer-associated biomarker proteins) (4) are making such an objective potentially achievable. These advanced technologies focus on a deeper understanding of plasma for the sensitive, specific and accurate measurement of early-stage cancer biomarkers.

However, historically most cancer-associated protein biomarkers have been reported in patient plasma at particularly low abundance. Indeed, this may be more problematic for early stage cancer detection when the tumour size is small, whilst the physiological and immune response to the cancer is minimal and cancer-associated biomarkers (shed or leaked proteins) will also be of low abundance. In the case of protein/peptide identification, this challenge is exacerbated by the high dynamic concentration range of proteins found in plasma, making identification of low abundance proteins biomarkers a daunting task. Most studies attempting to uncover cancer-associated plasma biomarkers use depletion, multidimensional fractionation or some form of enrichment (4, 5, 7, 33). Although MS-based technologies have been suggested to be more specific and accurate than antibody-based techniques, and are amenable to multiplex analysis, they are not yet compatible with high-throughput methodologies <u>and</u> often fail to identify very low abundance plasma proteins. Conversely, antibody-based technologies suffer from batch variation issues, non-specific binding (94) as well as cost and reliability of developing multiplex assays. In this study, we aimed to identify very low abundance cancer-associated proteins (see Table 1 for details) from non-depleted CRC plasma, including several not detectable by current MS methodologies, using a rPSL SWATH approach.

Although SWATH-MS has been available for protein analysis and quantification for a decade, the fundamental hurdle of extracting consistent, useful and reliable data from DIA runs remains. It should be noted that here we have <u>initially</u> deliberately focussed on protein identification rather than protein quantification. We applied the simple premise that if plasma is a repository of molecules reflecting the biological and physiological status of the human body (95), then DIA MS (here SWATH) should be able to detect all spectra (96) including low abundance protein biomarkers. Although there are a number of ways to investigate this, including multiple library free approaches (97), *in silico* spectral libraries (98), labelling (99), spiking (100) or synthetic peptide approaches (101), we chose to use a library approach using a recombinant protein library of 36 low-medium abundance since it was hoped that a more comprehensive tryptic peptide representation for each protein could be achieved.

Using a rPSL prior to SWATH on patient samples, we achieved an average of 70% coverage for each protein using DDA analysis, a level of coverage that would not be feasible (at a reasonable cost) with a synthetic peptide library or using DDA methods on similar biological plasma samples. This makes the recombinant protein DDA library approach more comprehensive for identifying low abundance proteins in a SWATH experiment. Additionally, routine, established and accepted pipelines for data analysis (14, 102) could be applied leading to more reliable results. In any SWATH-MS experiment, the more comprehensive the library (resolution, peptide number, protein coverage) the more accurate will be the identifications. To achieve a comprehensive library, we ran the recombinant proteins in discrete groups based on molecular weight groupings (Fig 1) to allow suitable coverage without peptides from smaller proteins overwhelming those from larger proteins. This allowed the identification of a high number of peptides for each protein in the rPSL, illustrating an accurate and comprehensive DDA library.

Thirty-two cancer-associated plasma proteins were detected using rPSL SWATH (from 36), of which 20 were exclusively detected when compared to a DDA plasma protein spectral library (Fig 2). Importantly, the reported plasma concentration of these proteins were below 10ng/ml, and 7 proteins were actually in the pg/ml range. Furthermore, 33 proteins were reliably identified from a merged library (rPSL + MARS-14 depleted plasma protein DDA spectral library) after the application of high stringency protein identification criteria, HPP Guideline v3.0 (Table 3) (29). The ability to reliably detect lower abundance disease-related plasma proteins has previously only been achieved with multidimensional fractionation, selective monitoring (or enrichment) or affinity-based approaches.

The rPSL SWATH allowed the identification of well-documented and clinically significant (supported by the literature) plasma proteins. Numerous studies have proposed CEA as a late-stage CRC biomarker (35), as well as for lung (36) and gastric cancers (37). Plasma IL6, CXCL8 and ILB1 have been also proposed as diagnostic markers in CRC, brain, breast, oral and prostate cancers (45–47, 55–57). Reported plasma concentrations of ILB1 and IL6 are both ~1ng/ml, and for CXCL8 are ~14pg/ml. Our study is, we believe, the first to show detection of these cytokines in human disease plasmas using MS. Furthermore, rPSL SWATH was able to identify CRC biomarkers ITGB6 and PLAUR implicated in our own previous studies (4, 23–25, 103). The reported plasma concentrations of both ITGB6 and PLAUR are ~2ng/ml and again this study is, to our knowledge, the first to demonstrate plasma ITGB6 detection using MS. PLAUR expression in tumour tissues and plasma has been recognised as a potential biomarker for many cancer types, including CRC (104). Notably, PLAUR measurement on tumour cell surfaces and detection of soluble-uPAR (suPAR) (cleaved uPAR isoforms released from the cell surface containing domains D1, D2+D3 or D1+D2+D3) in plasma have been recognised as prognostic indicators of CRC survival (70, 105). However, these studies used antibody-based technologies, and has issues with both non-specific binding and batch variation (94). We contend that developing high-throughput targeted MS prognostic tools employing a novel rPSL approach will be of significant benefit across many clinical settings. Collectively, identification and quantification of low abundance cancer-associated proteins from non-depleted or non-fractionated plasma allows a more seamless transition to potential clinical applications (100), eliminates the risk of depletion of unintended non-targeted proteins (106) and serves to demonstrate the efficiency of SWATH-MS.

Although our novel rPSL SWATH approach has substantial benefits for proteomics applications, there are still issues to be considered around FDR corrections when merging differently sized DDA libraries. In this study, we addressed the FDR concerns due to the use of a small library by merging two libraries (rPSL and plasma protein DDA library). SWATH analysis using this merged library resulted in three additional plasma protein identifications (ADAMDEC1, CST3 and PLAU). However, detected quantifiable peptides for these three proteins were derived from the rPSL part of the merge, not from the MARS-14 depleted plasma protein DDA library. We recognise that the rPSL is a relatively small library constructed using only 36 recombinant proteins and 1,435 peptide identifications. SWATH analysis at 1% FDR provided 32 protein identifications supported by 134 (or 9.3%) quantifiable peptides from non-depleted CRC plasma. In contrast, the merged library contained 762 proteins and 33,295 peptides, and SWATH analysis (1% FDA) resulted in 519 protein identifications with 3,187 (or 9.5%) quantifiable peptides. Although the ratio of quantifiable peptides between the rPSL (smaller) and merged library (larger) was similar, the actual peptide content was different due to the application of different FDR requirements for the different libraries, as well as the different library size. We assumed that, in rPSL, the peptides (for ADAMDEC1, CST3 and PLAU) may have been eliminated as false positives (i.e., not passed at the 1% FDR cut off for at least three replicates). However, when the libraries were merged, the number of false positive peptides from the larger library may have been overwhelmed such that eliminated peptides in rPSL may have been accepted as true positives in the larger merged library.

DDA experiments often suffer from stochastic selection of precursor ions for MS/MS fragmentation (84), and this especially applies to clinical plasma studies due to the complexity of the samples used. SWATH/DIA studies involving DDA library building also suffer similar stochastic selection issues. In the case of studies where a specific panel of proteins needs to be monitored routinely in a multiplexed manner, rPSL may be a suitable method as the rPSL library needs only to be developed once, and unlike a predicted library (as done by PROSIT (22)), these libraries have real empirical evidence. Hence, rPSL can be tailored to the specific protein detection panel, and once built can be permanently used. Thus, panels of hundreds, or even thousands, of proteins and resultant tryptic peptides can be detected simultaneously in a high-throughput manner.

In conclusion, this study used an rPSL SWATH approach to identify lower abundance cancer-associated proteins from non-depleted CRC plasmas. We were able to demonstrate the premise that this novel approach can probe deeper into the plasma proteome compared to standard DDA shotgun proteomics or DIA/SWATH where DDA libraries are generated from biological samples. Using rPSL SWATH, we were able to see 8 additional proteins that have never previously been observed by MS at high-stringency protein inference. The implication of this study is that MS technologies can reliably achieve picogram detection on non-depleted plasma (currently thought to be obtainable only using more sensitive antibody-based affinity methods). We contend that rPSL SWATH approaches can allow accurate, reliable and readily adaptable clinical measurement of multiple low abundance plasma biomarkers (panels) simultaneously in a single workflow.

## Supporting information

Skyline peptide and transition settings.

Identified proteins/peptides from DDA recombinant protein spectral library (rPSL).

Identified proteins/peptides from rPSL SWATH analysis - PeakView and Skyline.

Identified proteins/peptides from human plasma DDA library.

Identified proteins/peptides from merged library SWATH analysis.

A list of identified peptides from rPSL SWATH and merged library SWATH.

Peptides (for ADAMDEC1, CST3 and PLAU) identified from rPSL, human plasma DDA library and merged library SWATH analysis.

Identified proteins and peptides from merged library SWATH - before and after applying high stringency protein identification criteria.

## ASSOCIATED CONTENT

## Supporting information

Supplementary Table S1: Skyline peptide and transition settings. Supplementary Table S2: Identified proteins/peptides from DDA recombinant protein spectral library (rPSL). Supplementary Table S3: Identified proteins/peptides from rPSL SWATH analysis - PeakView and Skyline. Supplementary Table S4: Identified proteins/peptides from human plasma DDA library. Supplementary Table S5: Identified proteins/peptides from merged library SWATH analysis. Supplementary Table S6: A list of identified peptides from rPSL SWATH and merged library SWATH. Supplementary Table S7: Peptides (for ADAMDEC1, CST3 and PLAU) identified from rPSL, human plasma DDA library and merged library SWATH analysis. Supplementary Table S8: Identified proteins and peptides from merged library SWATH - before and after applying high stringency protein identification criteria. Mass spectrometry data is available through the ProteomeXchange consortium via the PRIDE partner repository with the dataset identifier PXD022361.

## AUTHOR INFORMATION

## Corresponding Authors

**Seong Beom Ahn** – Department of Biomedical Sciences, Faculty of Medicine and Health Sciences, Macquarie University, NSW, 2109, Australia;

**Mark S. Baker** – Department of Biomedical Sciences, Faculty of Medicine and Health Sciences, Macquarie University, NSW, 2109, Australia;

## Authors

**Karthik Shantharam Kamath** – Australian Proteome Analysis Facility (APAF), Department of Molecular Sciences, Faculty of Science and Engineering, Macquarie University, NSW, 2109, Australia.

**Abidali Mohamedali** – Department of Molecular Sciences, Faculty of Science and Engineering, Macquarie University, NSW, 2109, Australia.

**Zainab Noor** – ProCan^®^, Children’s Medical Research Institute, The University of Sydney, Westmead, NSW, Australia.

**Jemma X. Wu** – Australian Proteome Analysis Facility (APAF), Department of Molecular Sciences, Faculty of Science and Engineering, Macquarie University, NSW, 2109, Australia.

**Dana Pascovici** – Australian Proteome Analysis Facility (APAF), Department of Molecular Sciences, Faculty of Science and Engineering, Macquarie University, NSW, 2109, Australia.

**Subash Adhikari** – Department of Biomedical Sciences, Faculty of Medicine and Health Sciences, Macquarie University, NSW, 2109, Australia.

**Harish R Cheruku** – Department of Biomedical Sciences, Faculty of Medicine and Health Sciences, Macquarie University, NSW, 2109, Australia.

**Gilles J. Guillemin** – Department of Biomedical Sciences, Faculty of Medicine and Health Sciences, Macquarie University, NSW, 2109, Australia.

**Matthew J. McKay** – Australian Proteome Analysis Facility (APAF), Department of Molecular Sciences, Faculty of Science and Engineering, Macquarie University, NSW, 2109, Australia.

**Edouard C. Nice** – Department of Biochemistry and Molecular Biology, Faculty of Medicine, Nursing and Health Sciences, Monash University, VIC, 3800, Australia.

## Author Contributions

SBA, MSB designed experiments. SBA, KSK, AM, SA, MJM performed experiments. SBA, KSK, AM, ZN, JXW, DP performed MS data and statistical analyses. SBA, MSB, KSK, AM, SA, JXW, DP, HRC, GJG, ECN prepared figures and tables. All authors contributed to writing/reviewing of each manuscript version.

## ACKNOWLEDGMENTS

The authors acknowledge and thank to the Victorian Cancer Biobank for providing CRC patient EDTA-plasma samples. This study was supported by the Cancer Institute NSW ECR fellowship 15/ECF/1-38 (SBA), Cancer Council NSW RG19-04 (MSB, SBA, ECN), NHMRC project grant 1010303 (MSB, ECN), ‘Fight on the Beaches’ (MSB, SBA, ECN, SA), Sydney Vital CINSW Translational Cancer Research Centre grant (MSB, SBA, SA) and iMQRES funding from Macquarie University (SA).

## References

1. Dakubo, G. D., Advanced Technologies for Body Fluid Biomarker Analyses. In Cancer Biomarkers in Body Fluids, Springer: 2016; pp 55–74.

2. Ridker, P. M.; Rifai, N.; Stampfer, M. J.; Hennekens, C. H., Plasma concentration of interleukin-6 and the risk of future myocardial infarction among apparently healthy men. Circulation 2000, 101, (15), 1767–72.

3. Geyer, P. E.; Holdt, L. M.; Teupser, D.; Mann, M., Revisiting biomarker discovery by plasma proteomics. Mol Syst Biol 2017, 13, (9), 942.

4. Ahn, S. B.; Sharma, S.; Mohamedali, A.; Mahboob, S.; Redmond, W. J.; Pascovici, D.; Wu, J. X.; Zaw, T.; Adhikari, S.; Vaibhav, V.; Nice, E. C.; Baker, M. S., Potential early clinical stage colorectal cancer diagnosis using a proteomics blood test panel. Clin Proteomics 2019, 16, 34.

5. Yadav, A. K.; Bhardwaj, G.; Basak, T.; Kumar, D.; Ahmad, S.; Priyadarshini, R.; Singh, A. K.; Dash, D.; Sengupta, S., A systematic analysis of eluted fraction of plasma post immunoaffinity depletion: implications in biomarker discovery. PLoS One 2011, 6, (9), e24442.

6. Nice, E. C.; Rothacker, J.; Weinstock, J.; Lim, L.; Catimel, B., Use of multidimensional separation protocols for the purification of trace components in complex biological samples for proteomics analysis. J Chromatogr A 2007, 1168, (1-2), 190–210; discussion 189.

7. Seong, Y.; Yoo, Y. S.; Akter, H.; Kang, M. J., Sample preparation for detection of low abundance proteins in human plasma using ultra-high performance liquid chromatography coupled with highly accurate mass spectrometry. J Chromatogr B Analyt Technol Biomed Life Sci 2017, 1060, 272–280.

8. Ahn, S.-B.; Khan, A., Detection and quantitation of twenty-seven cytokines, chemokines and growth factors pre- and post-high abundance protein depletion in human plasma. EuPA Open Proteomics 2014, 3, 78–84.

9. Lim, S. Y.; Lee, J. H.; Welsh, S. J.; Ahn, S. B.; Breen, E.; Khan, A.; Carlino, M. S.; Menzies, A. M.; Kefford, R. F.; Scolyer, R. A.; Long, G. V.; Rizos, H., Evaluation of two high-throughput proteomic technologies for plasma biomarker discovery in immunotherapy-treated melanoma patients. Biomark Res 2017, 5, 32.

10. Tu, C.; Rudnick, P. A.; Martinez, M. Y.; Cheek, K. L.; Stein, S. E.; Slebos, R. J.; Liebler, D. C., Depletion of abundant plasma proteins and limitations of plasma proteomics. J Proteome Res 2010, 9, (10), 4982–91.

11. Keshishian, H.; Burgess, M. W.; Gillette, M. A.; Mertins, P.; Clauser, K. R.; Mani, D. R.; Kuhn, E. W.; Farrell, L. A.; Gerszten, R. E.; Carr, S. A., Multiplexed, Quantitative Workflow for Sensitive Biomarker Discovery in Plasma Yields Novel Candidates for Early Myocardial Injury. Mol Cell Proteomics 2015, 14, (9), 2375–93.

12. Geyer, P. E.; Kulak, N. A.; Pichler, G.; Holdt, L. M.; Teupser, D.; Mann, M., Plasma Proteome Profiling to Assess Human Health and Disease. Cell Syst 2016, 2, (3), 185–95.

13. Schwenk, J. M.; Omenn, G. S.; Sun, Z.; Campbell, D. S.; Baker, M. S.; Overall, C. M.; Aebersold, R.; Moritz, R. L.; Deutsch, E. W., The Human Plasma Proteome Draft of 2017: Building on the Human Plasma PeptideAtlas from Mass Spectrometry and Complementary Assays. J Proteome Res 2017, 16, (12), 4299–4310.

14. Ludwig, C.; Gillet, L.; Rosenberger, G.; Amon, S.; Collins, B. C.; Aebersold, R., Data-independent acquisition-based SWATH-MS for quantitative proteomics: a tutorial. Mol Syst Biol 2018, 14, (8), e8126.

15. Wu, J. X.; Song, X.; Pascovici, D.; Zaw, T.; Care, N.; Krisp, C.; Molloy, M. P., SWATH Mass Spectrometry Performance Using Extended Peptide MS/MS Assay Libraries. Mol Cell Proteomics 2016, 15, (7), 2501–14.

16. Schubert, O. T.; Gillet, L. C.; Collins, B. C.; Navarro, P.; Rosenberger, G.; Wolski, W. E.; Lam, H.; Amodei, D.; Mallick, P.; MacLean, B.; Aebersold, R., Building high-quality assay libraries for targeted analysis of SWATH MS data. Nat Protoc 2015, 10, (3), 426–41.

17. Gillet, L. C.; Navarro, P.; Tate, S.; Rost, H.; Selevsek, N.; Reiter, L.; Bonner, R.; Aebersold, R., Targeted data extraction of the MS/MS spectra generated by data-independent acquisition: a new concept for consistent and accurate proteome analysis. Mol Cell Proteomics 2012, 11, (6), O111 016717.

18. Noor, Z.; Ahn, S. B.; Baker, M. S.; Ranganathan, S.; Mohamedali, A., Mass spectrometry-based protein identification in proteomics-a review. Brief Bioinform 2020.

19. Zi, J.; Zhang, S.; Zhou, R.; Zhou, B.; Xu, S.; Hou, G.; Tan, F.; Wen, B.; Wang, Q.; Lin, L.; Liu, S., Expansion of the ion library for mining SWATH-MS data through fractionation proteomics. Anal Chem 2014, 86, (15), 7242–6.

20. Rosenberger, G.; Koh, C. C.; Guo, T.; Rost, H. L.; Kouvonen, P.; Collins, B. C.; Heusel, M.; Liu, Y.; Caron, E.; Vichalkovski, A.; Faini, M.; Schubert, O. T.; Faridi, P.; Ebhardt, H. A.; Matondo, M.; Lam, H.; Bader, S. L.; Campbell, D. S.; Deutsch, E. W.; Moritz, R. L.; Tate, S.; Aebersold, R., A repository of assays to quantify 10,000 human proteins by SWATH-MS. Sci Data 2014, 1, 140031.

21. Wu, J. X.; Pascovici, D.; Ignjatovic, V.; Song, X.; Krisp, C.; Molloy, M. P., Improving Protein Detection Confidence Using SWATH-Mass Spectrometry with Large Peptide Reference Libraries. Proteomics 2017, 17, (19).

22. Gessulat, S.; Schmidt, T.; Zolg, D. P.; Samaras, P.; Schnatbaum, K.; Zerweck, J.; Knaute, T.; Rechenberger, J.; Delanghe, B.; Huhmer, A.; Reimer, U.; Ehrlich, H. C.; Aiche, S.; Kuster, B.; Wilhelm, M., Prosit: proteome-wide prediction of peptide tandem mass spectra by deep learning. Nat Methods 2019, 16, (6), 509–518.

23. Ahn, S. B.; Chan, C.; Dent, O. F.; Mohamedali, A.; Kwun, S. Y.; Clarke, C.; Fletcher, J.; Chapuis, P. H.; Nice, E. C.; Baker, M. S., Epithelial and stromal cell urokinase plasminogen activator receptor expression differentially correlates with survival in rectal cancer stages B and C patients. PLoS One 2015, 10, (2), e0117786.

24. Ahn, S. B.; Mohamedali, A.; Chan, C.; Fletcher, J.; Kwun, S. Y.; Clarke, C.; Dent, O. F.; Chapuis, P. H.; Nice, E.; Baker, M. S., Correlations between integrin ανβ6 expression and clinico-pathological features in stage B and stage C rectal cancer. PLoS One 2014, 9, (5), e97248.

25. Ahn, S. B.; Mohamedali, A.; Pascovici, D.; Adhikari, S.; Sharma, S.; Nice, E. C.; Baker, M. S., Proteomics Reveals Cell-Surface Urokinase Plasminogen Activator Receptor Expression Impacts Most Hallmarks of Cancer. Proteomics 2019, 19, (21–22), e1900026.

26. Mahboob, S.; Ahn, S. B.; Cheruku, H. R.; Cantor, D.; Rennel, E.; Fredriksson, S.; Edfeldt, G.; Breen, E. J.; Khan, A.; Mohamedali, A.; Muktadir, M. G.; Ranganathan, S.; Tan, S. H.; Nice, E.; Baker, M. S., A novel multiplexed immunoassay identifies CEA, IL-8 and prolactin as prospective markers for Dukes’ stages A-D colorectal cancers. Clin Proteomics 2015, 12, (1), 10.

27. Deutsch, E. W.; Lane, L.; Overall, C. M.; Bandeira, N.; Baker, M. S.; Pineau, C.; Moritz, R. L.; Corrales, F.; Orchard, S.; Van Eyk, J. E.; Paik, Y. K.; Weintraub, S. T.; Vandenbrouck, Y.; Omenn, G. S., Human Proteome Project Mass Spectrometry Data Interpretation Guidelines 3.0. J Proteome Res 2019, 18, (12), 4108–4116.

28. Adhikari, S.; Nice, E. C.; Deutsch, E. W.; Lane, L.; Omenn, G. S.; Pennington, S. R.; Paik, Y. K.; Overall, C. M.; Corrales, F. J.; Cristea, I. M.; Van Eyk, J. E.; Uhlén, M.; Lindskog, C.; Chan, D. W.; Bairoch, A.; Waddington, J. C.; Justice, J. L.; LaBaer, J.; Rodriguez, H.; He, F.; Kostrzewa, M.; Ping, P.; Gundry, R. L.; Stewart, P.; Srivastava, S.; Srivastava, S.; Nogueira, F. C. S.; Domont, G. B.; Vandenbrouck, Y.; Lam, M. P. Y.; Wennersten, S.; Vizcaino, J. A.; Wilkins, M.; Schwenk, J. M.; Lundberg, E.; Bandeira, N.; Marko-Varga, G.; Weintraub, S. T.; Pineau, C.; Kusebauch, U.; Moritz, R. L.; Ahn, S. B.; Palmblad, M.; Snyder, M. P.; Aebersold, R.; Baker, M. S., A high-stringency blueprint of the human proteome. Nat Commun 2020, 11, (1), 5301.

29. Liu, T.; Qian, W. J.; Gritsenko, M. A.; Xiao, W.; Moldawer, L. L.; Kaushal, A.; Monroe, M. E.; Varnum, S. M.; Moore, R. J.; Purvine, S. O.; Maier, R. V.; Davis, R. W.; Tompkins, R. G.; Camp, D. G., 2nd; Smith, R. D.; Inflammation; the Host Response to Injury Large Scale Collaborative Research, P., High dynamic range characterization of the trauma patient plasma proteome. Mol Cell Proteomics 2006, 5, (10), 1899–913.

30. Kume, H.; Muraoka, S.; Kuga, T.; Adachi, J.; Narumi, R.; Watanabe, S.; Kuwano, M.; Kodera, Y.; Matsushita, K.; Fukuoka, J.; Masuda, T.; Ishihama, Y.; Matsubara, H.; Nomura, F.; Tomonaga, T., Discovery of colorectal cancer biomarker candidates by membrane proteomic analysis and subsequent verification using selected reaction monitoring (SRM) and tissue microarray (TMA) analysis. Mol Cell Proteomics 2014, 13, (6), 1471–84.

31. Jiang, W.; Huang, R.; Duan, C.; Fu, L.; Xi, Y.; Yang, Y.; Yang, W. M.; Yang, D.; Yang, D. H.; Huang, R. P., Identification of five serum protein markers for detection of ovarian cancer by antibody arrays. PLoS One 2013, 8, (10), e76795.

32. Ueda, K.; Tatsuguchi, A.; Saichi, N.; Toyama, A.; Tamura, K.; Furihata, M.; Takata, R.; Akamatsu, S.; Igarashi, M.; Nakayama, M.; Sato, T. A.; Ogawa, O.; Fujioka, T.; Shuin, T.; Nakamura, Y.; Nakagawa, H., Plasma low-molecular-weight proteome profiling identified neuropeptide-Y as a prostate cancer biomarker polypeptide. J Proteome Res 2013, 12, (10), 4497–506.

33. Whiteaker, J. R.; Zhang, H.; Eng, J. K.; Fang, R.; Piening, B. D.; Feng, L. C.; Lorentzen, T. D.; Schoenherr, R. M.; Keane, J. F.; Holzman, T.; Fitzgibbon, M.; Lin, C.; Zhang, H.; Cooke, K.; Liu, T.; Camp, D. G., 2nd; Anderson, L.; Watts, J.; Smith, R. D.; McIntosh, M. W.; Paulovich, A. G., Head-to-head comparison of serum fractionation techniques. J Proteome Res 2007, 6, (2), 828–36.

34. Yan, B.; Chen, B.; Min, S.; Gao, Y.; Zhang, Y.; Xu, P.; Li, C.; Chen, J.; Luo, G.; Liu, C., iTRAQ-based Comparative Serum Proteomic Analysis of Prostate Cancer Patients with or without Bone Metastasis. J Cancer 2019, 10, (18), 4165–4177.

35. Christensen, I. J.; Brunner, N.; Dowell, B.; Davis, G.; Nielsen, H. J.; Newstead, G.; King, D., Plasma TIMP-1 and CEA as Markers for Detection of Primary Colorectal Cancer: A Prospective Validation Study Including Symptomatic and Non-symptomatic Individuals. Anticancer Res 2015, 35, (9), 4935–41.

36. Djureinovic, D.; Ponten, V.; Landelius, P.; Al Sayegh, S.; Kappert, K.; Kamali-Moghaddam, M.; Micke, P.; Stahle, E., Multiplex plasma protein profiling identifies novel markers to discriminate patients with adenocarcinoma of the lung. BMC Cancer 2019, 19, (1), 741.

37. Shen, Q.; Polom, K.; Williams, C.; de Oliveira, F. M. S.; Guergova-Kuras, M.; Lisacek, F.; Karlsson, N. G.; Roviello, F.; Kamali-Moghaddam, M., A targeted proteomics approach reveals a serum protein signature as diagnostic biomarker for resectable gastric cancer. EBioMedicine 2019, 44, 322–333.

38. Braga-Lagache, S.; Buchs, N.; Iacovache, M. I.; Zuber, B.; Jackson, C. B.; Heller, M., Robust Label-free, Quantitative Profiling of Circulating Plasma Microparticle (MP) Associated Proteins. Mol Cell Proteomics 2016, 15, (12), 3640–3652.

39. Geyer, P. E.; Wewer Albrechtsen, N. J.; Tyanova, S.; Grassl, N.; Iepsen, E. W.; Lundgren, J.; Madsbad, S.; Holst, J. J.; Torekov, S. S.; Mann, M., Proteomics reveals the effects of sustained weight loss on the human plasma proteome. Mol Syst Biol 2016, 12, (12), 901.

40. Lee, J. H.; Cho, H. S.; Lee, J. J.; Jun, S. Y.; Ahn, J. H.; Min, J. S.; Yoon, J. Y.; Choi, M. H.; Jeon, S. J.; Lim, J. H.; Jung, C. R.; Kim, D. S.; Kim, H. T.; Factor, V. M.; Lee, Y. H.; Thorgeirsson, S. S.; Kim, C. H.; Kim, N. S., Plasma glutamate carboxypeptidase is a negative regulator in liver cancer metastasis. Oncotarget 2016, 7, (48), 79774–79786.

41. Chen, R.; Mias, G. I.; Li-Pook-Than, J.; Jiang, L.; Lam, H. Y.; Chen, R.; Miriami, E.; Karczewski, K. J.; Hariharan, M.; Dewey, F. E.; Cheng, Y.; Clark, M. J.; Im, H.; Habegger, L.; Balasubramanian, S.; O’Huallachain, M.; Dudley, J. T.; Hillenmeyer, S.; Haraksingh, R.; Sharon, D.; Euskirchen, G.; Lacroute, P.; Bettinger, K.; Boyle, A. P.; Kasowski, M.; Grubert, F.; Seki, S.; Garcia, M.; Whirl-Carrillo, M.; Gallardo, M.; Blasco, M. A.; Greenberg, P. L.; Snyder, P.; Klein, T. E.; Altman, R. B.; Butte, A. J.; Ashley, E. A.; Gerstein, M.; Nadeau, K. C.; Tang, H.; Snyder, M., Personal omics profiling reveals dynamic molecular and medical phenotypes. Cell 2012, 148, (6), 1293–307.

42. Greening, D. W.; Simpson, R. J., A centrifugal ultrafiltration strategy for isolating the low-molecular weight (<or=25K) component of human plasma proteome. J Proteomics 2010, 73, (3), 637–48.

43. Bolke, E.; Schieren, G.; Gripp, S.; Steinbach, G.; Peiper, M.; Orth, K.; Matuschek, C.; Pelzer, M.; Lammering, G.; Houben, R.; Antke, C.; Rump, L. C.; Mota, R.; Gerber, P. A.; Schuler, P.; Hoffmann, T. K.; Rusnak, E.; Hermsen, D.; Budach, W., Cystatin C - a fast and reliable biomarker for glomerular filtration rate in head and neck cancer patients. Strahlenther Onkol 2011, 187, (3), 191–201.

44. Edwards, N. J.; Oberti, M.; Thangudu, R. R.; Cai, S.; McGarvey, P. B.; Jacob, S.; Madhavan, S.; Ketchum, K. A., The CPTAC Data Portal: A Resource for Cancer Proteomics Research. J Proteome Res 2015, 14, (6), 2707–13.

45. Bunger, S.; Haug, U.; Kelly, F. M.; Klempt-Giessing, K.; Cartwright, A.; Posorski, N.; Dibbelt, L.; Fitzgerald, S. P.; Bruch, H. P.; Roblick, U. J.; von Eggeling, F.; Brenner, H.; Habermann, J. K.; Chip, B. M.-C. C. C. S., Toward standardized high-throughput serum diagnostics: multiplex-protein array identifies IL-8 and VEGF as serum markers for colon cancer. J Biomol Screen 2011, 16, (9), 1018–26.

46. Ilhan-Mutlu, A.; Wagner, L.; Widhalm, G.; Wohrer, A.; Bartsch, S.; Czech, T.; Heinzl, H.; Leutmezer, F.; Prayer, D.; Marosi, C.; Base, W.; Preusser, M., Exploratory investigation of eight circulating plasma markers in brain tumor patients. Neurosurg Rev 2013, 36, (1), 45–55; discussion 55–6.

47. Todorovic-Rakovic, N.; Milovanovic, J., Interleukin-8 in breast cancer progression. J Interferon Cytokine Res 2013, 33, (10), 563–70.

48. Narita, D.; Seclaman, E.; Anghel, A.; Ilina, R.; Cireap, N.; Negru, S.; Sirbu, I. O.; Ursoniu, S.; Marian, C., Altered levels of plasma chemokines in breast cancer and their association with clinical and pathological characteristics. Neoplasma 2016, 63, (1), 141–9.

49. Guldbrandsen, A.; Vethe, H.; Farag, Y.; Oveland, E.; Garberg, H.; Berle, M.; Myhr, K. M.; Opsahl, J. A.; Barsnes, H.; Berven, F. S., In-depth characterization of the cerebrospinal fluid (CSF) proteome displayed through the CSF proteome resource (CSF-PR). Mol Cell Proteomics 2014, 13, (11), 3152–63.

50. Lavaee, F.; Zare, S.; Mojtahedi, Z.; Malekzadeh, M.; Khademi, B.; Ghaderi, A., Serum CXCL12, but not CXCR4, Is Associated with Head and Neck Squamous Cell Carcinomas. Asian Pac J Cancer Prev 2018, 19, (4), 901–904.

51. Lukaszewicz-Zajac, M.; Mroczko, B.; Kozlowski, M.; Szmitkowski, M., The Serum Concentrations of Chemokine CXCL12 and Its Specific Receptor CXCR4 in Patients with Esophageal Cancer. Dis Markers 2016, 2016, 7963895.

52. Blanco-Prieto, S.; Vazquez-Iglesias, L.; Rodriguez-Girondo, M.; Barcia-Castro, L.; Fernandez-Villar, A.; Botana-Rial, M. I.; Rodriguez-Berrocal, F. J.; de la Cadena, M. P., Serum calprotectin, CD26 and EGF to establish a panel for the diagnosis of lung cancer. PLoS One 2015, 10, (5), e0127318.

53. Omenn, G. S.; States, D. J.; Adamski, M.; Blackwell, T. W.; Menon, R.; Hermjakob, H.; Apweiler, R.; Haab, B. B.; Simpson, R. J.; Eddes, J. S.; Kapp, E. A.; Moritz, R. L.; Chan, D. W.; Rai, A. J.; Admon, A.; Aebersold, R.; Eng, J.; Hancock, W. S.; Hefta, S. A.; Meyer, H.; Paik, Y. K.; Yoo, J. S.; Ping, P.; Pounds, J.; Adkins, J.; Qian, X.; Wang, R.; Wasinger, V.; Wu, C. Y.; Zhao, X.; Zeng, R.; Archakov, A.; Tsugita, A.; Beer, I.; Pandey, A.; Pisano, M.; Andrews, P.; Tammen, H.; Speicher, D. W.; Hanash, S. M., Overview of the HUPO Plasma Proteome Project: results from the pilot phase with 35 collaborating laboratories and multiple analytical groups, generating a core dataset of 3020 proteins and a publicly-available database. Proteomics 2005, 5, (13), 3226–45.

54. Banys-Paluchowski, M.; Witzel, I.; Riethdorf, S.; Rack, B.; Janni, W.; Fasching, P. A.; Solomayer, E. F.; Aktas, B.; Kasimir-Bauer, S.; Pantel, K.; Fehm, T.; Muller, V., Evaluation of serum epidermal growth factor receptor (EGFR) in correlation to circulating tumor cells in patients with metastatic breast cancer. Sci Rep 2017, 7, (1), 17307.

55. Brailo, V.; Vucicevic-Boras, V.; Lukac, J.; Biocina-Lukenda, D.; Zilic-Alajbeg, I.; Milenovic, A.; Balija, M., Salivary and serum interleukin 1 beta, interleukin 6 and tumor necrosis factor alpha in patients with leukoplakia and oral cancer. Med Oral Patol Oral Cir Bucal 2012, 17, (1), e10–5.

56. Lu, C. C.; Kuo, H. C.; Wang, F. S.; Jou, M. H.; Lee, K. C.; Chuang, J. H., Upregulation of TLRs and IL-6 as a marker in human colorectal cancer. Int J Mol Sci 2014, 16, (1), 159–77.

57. Shariat, S. F.; Semjonow, A.; Lilja, H.; Savage, C.; Vickers, A. J.; Bjartell, A., Tumor markers in prostate cancer I: blood-based markers. Acta Oncol 2011, 50 Suppl 1, 61–75.

58. Ahmed, N.; Riley, C.; Rice, G. E.; Quinn, M. A.; Baker, M. S., Alpha(v)beta(6) integrin-A marker for the malignant potential of epithelial ovarian cancer. J Histochem Cytochem 2002, 50, (10), 1371–80.

59. dos Santos, P. B.; Zanetti, J. S.; Ribeiro-Silva, A.; Beltrao, E. I., Beta 1 integrin predicts survival in breast cancer: a clinicopathological and immunohistochemical study. Diagn Pathol 2012, 7, 104.

60. Bengs, S.; Becker, E.; Busenhart, P.; Spalinger, M. R.; Raselli, T.; Kasper, S.; Lang, S.; Atrott, K.; Mamie, C.; Vavricka, S. R.; von Boehmer, L.; Knuth, A.; Tuomisto, A.; Makinen, M. J.; Hruz, P.; Turina, M.; Rickenbacher, A.; Petrowsky, H.; Weber, A.; Frei, P.; Halama, M.; Jenkins, G.; Sheppard, D.; Croner, R. S.; Christoph, J.; Britzen-Laurent, N.; Naschberger, E.; Schellerer, V.; Sturzl, M.; Fried, M.; Rogler, G.; Scharl, M., beta6-integrin serves as a novel serum tumor marker for colorectal carcinoma. Int J Cancer 2019, 145, (3), 678–685.

61. Larkin, S. E.; Johnston, H. E.; Jackson, T. R.; Jamieson, D. G.; Roumeliotis, T. I.; Mockridge, C. I.; Michael, A.; Manousopoulou, A.; Papachristou, E. K.; Brown, M. D.; Clarke, N. W.; Pandha, H.; Aukim-Hastie, C. L.; Cragg, M. S.; Garbis, S. D.; Townsend, P. A., Detection of candidate biomarkers of prostate cancer progression in serum: a depletion-free 3D LC/MS quantitative proteomics pilot study. Br J Cancer 2016, 115, (9), 1078–1086.

62. Kluger, H. M.; Hoyt, K.; Bacchiocchi, A.; Mayer, T.; Kirsch, J.; Kluger, Y.; Sznol, M.; Ariyan, S.; Molinaro, A.; Halaban, R., Plasma markers for identifying patients with metastatic melanoma. Clin Cancer Res 2011, 17, (8), 2417–25.

63. Helgeland, E.; Breivik, L. E.; Vaudel, M.; Svendsen, O. S.; Garberg, H.; Nordrehaug, J. E.; Berven, F. S.; Jonassen, A. K., Exploring the human plasma proteome for humoral mediators of remote ischemic preconditioning--a word of caution. PLoS One 2014, 9, (10), e109279.

64. Dragutinovic, V. V.; Radonjic, N. V.; Petronijevic, N. D.; Tatic, S. B.; Dimitrijevic, I. B.; Radovanovic, N. S.; Krivokapic, Z. V., Matrix metalloproteinase-2 (MMP-2) and −9 (MMP-9) in preoperative serum as independent prognostic markers in patients with colorectal cancer. Mol Cell Biochem 2011, 355, (1–2), 173–8.

65. Patel, S.; Sumitra, G.; Koner, B. C.; Saxena, A., Role of serum matrix metalloproteinase-2 and −9 to predict breast cancer progression. Clin Biochem 2011, 44, (10–11), 869–72.

66. Cymbaluk-Ploska, A.; Chudecka-Glaz, A.; Pius-Sadowska, E.; Machalinski, B.; Menkiszak, J.; Sompolska-Rzechula, A., Suitability assessment of baseline concentration of MMP3, TIMP3, HE4 and CA125 in the serum of patients with ovarian cancer. J Ovarian Res 2018, 11, (1), 1.

67. Zhong, X. Y.; Kaul, S.; Bastert, G., Evaluation of MUC1 and EGP40 in bone marrow and peripheral blood as a marker for occult breast cancer. Arch Gynecol Obstet 2001, 264, (4), 177–81.

68. Braicu, C.; Tudoran, O.; Balacescu, L.; Catana, C.; Neagoe, E.; Berindan-Neagoe, I.; Ionescu, C., The significance of PDGF expression in serum of colorectal carcinoma patients--correlation with Duke’s classification. Can PDGF become a potential biomarker? Chirurgia (Bucur) 2013, 108, (6), 849–54.

69. Shevchenko, V. E.; Kovalev, S. V.; Arnotskaya, N. E.; Zborovskaya, I. B.; Akhmedov, B. B.; Polotskii, B. E.; Kostin, A. U.; Moukeria, A. F.; Zaridze, D. G.; Davidov, M. I., Human blood plasma proteome mapping for search of potential markers of the lung squamous cell carcinoma. Eur J Mass Spectrom (Chichester) 2013, 19, (2), 123–33.

70. Lomholt, A. F.; Christensen, I. J.; Hoyer-Hansen, G.; Nielsen, H. J., Prognostic value of intact and cleaved forms of the urokinase plasminogen activator receptor in a retrospective study of 518 colorectal cancer patients. Acta Oncol 2010, 49, (6), 805–11.

71. Almasi, C. E.; Hoyer-Hansen, G.; Christensen, I. J.; Pappot, H., Prognostic significance of urokinase plasminogen activator receptor and its cleaved forms in blood from patients with non-small cell lung cancer. APMIS 2009, 117, (10), 755–61.

72. Kjellman, A.; Akre, O.; Gustafsson, O.; Hoyer-Hansen, G.; Lilja, H.; Norming, U.; Piironen, T.; Tornblom, M., Soluble urokinase plasminogen activator receptor as a prognostic marker in men participating in prostate cancer screening. J Intern Med 2011, 269, (3), 299–305.

73. Cheng, Y.; Liu, C.; Zhang, N.; Wang, S.; Zhang, Z., Proteomics analysis for finding serum markers of ovarian cancer. Biomed Res Int 2014, 2014, 179040.

74. Schmitz, M.; Grignard, G.; Margue, C.; Dippel, W.; Capesius, C.; Mossong, J.; Nathan, M.; Giacchi, S.; Scheiden, R.; Kieffer, N., Complete loss of PTEN expression as a possible early prognostic marker for prostate cancer metastasis. Int J Cancer 2007, 120, (6), 1284–92.

75. Minami, S.; Sato, Y.; Matsumoto, T.; Kageyama, T.; Kawashima, Y.; Yoshio, K.; Ishii, J.; Matsumoto, K.; Nagashio, R.; Okayasu, I., Proteomic study of sera from patients with bladder cancer: usefulness of S100A8 and S100A9 proteins. Cancer Genomics Proteomics 2010, 7, (4), 181–9.

76. Lawicki, S.; Zajkowska, M.; Glazewska, E. K.; Bedkowska, G. E.; Szmitkowski, M., Plasma levels and diagnostic utility of VEGF, MMP-9, and TIMP-1 in the diagnosis of patients with breast cancer. Onco Targets Ther 2016, 9, 911–9.

77. Lee, J. H.; Choi, J. W.; Kim, Y. S., Plasma or serum TIMP-1 is a predictor of survival outcomes in colorectal cancer: a meta-analysis. J Gastrointestin Liver Dis 2011, 20, (3), 287–91.

78. Mroczko, B.; Lukaszewicz-Zajac, M.; Gryko, M.; Kedra, B.; Szmitkowski, M., Clinical significance of serum levels of matrix metalloproteinase 2 (MMP-2) and its tissue inhibitor (TIMP-2) in gastric cancer. Folia Histochem Cytobiol 2011, 49, (1), 125–31.

79. Rieckmann, J. C.; Geiger, R.; Hornburg, D.; Wolf, T.; Kveler, K.; Jarrossay, D.; Sallusto, F.; Shen-Orr, S. S.; Lanzavecchia, A.; Mann, M.; Meissner, F., Social network architecture of human immune cells unveiled by quantitative proteomics. Nat Immunol 2017, 18, (5), 583–593.

80. Agnoli, C.; Grioni, S.; Pala, V.; Allione, A.; Matullo, G.; Gaetano, C. D.; Tagliabue, G.; Sieri, S.; Krogh, V., Biomarkers of inflammation and breast cancer risk: a case-control study nested in the EPIC-Varese cohort. Sci Rep 2017, 7, (1), 12708.

81. Chen, C.; Shen, H.; Zhang, L. G.; Liu, J.; Cao, X. G.; Yao, A. L.; Kang, S. S.; Gao, W. X.; Han, H.; Cao, F. H.; Li, Z. G., Construction and analysis of protein-protein interaction networks based on proteomics data of prostate cancer. Int J Mol Med 2016, 37, (6), 1576–86.

82. Dahse, R.; Utting, M.; Werner, W.; Schimmel, B.; Claussen, U.; Junker, K., TP53 alterations as a potential diagnostic marker in superficial bladder carcinoma and in patients serum, plasma and urine samples. International Journal of Oncology 2002, 20, (1), 107–115.

83. Nanjappa, V.; Thomas, J. K.; Marimuthu, A.; Muthusamy, B.; Radhakrishnan, A.; Sharma, R.; Ahmad Khan, A.; Balakrishnan, L.; Sahasrabuddhe, N. A.; Kumar, S.; Jhaveri, B. N.; Sheth, K. V.; Kumar Khatana, R.; Shaw, P. G.; Srikanth, S. M.; Mathur, P. P.; Shankar, S.; Nagaraja, D.; Christopher, R.; Mathivanan, S.; Raju, R.; Sirdeshmukh, R.; Chatterjee, A.; Simpson, R. J.; Harsha, H. C.; Pandey, A.; Prasad, T. S., Plasma Proteome Database as a resource for proteomics research: 2014 update. Nucleic Acids Res 2014, 42, (Database issue), D959–65.

84. Gillet, L. C.; Navarro, P.; Tate, S.; Röst, H.; Selevsek, N.; Reiter, L.; Bonner, R.; Aebersold, R., Targeted data extraction of the MS/MS spectra generated by data-independent acquisition: a new concept for consistent and accurate proteome analysis. Mol Cell Proteomics 2012, 11, (6), O111.016717.

85. Adhikari, S.; Sharma, S.; Ahn, S. B.; Baker, M. S., In Silico Peptide Repertoire of Human Olfactory Receptor Proteomes on High-Stringency Mass Spectrometry. J Proteome Res 2019, 18, (12), 4117–4123.

86. Schaeffer, M.; Gateau, A.; Teixeira, D.; Michel, P. A.; Zahn-Zabal, M.; Lane, L., The neXtProt peptide uniqueness checker: a tool for the proteomics community. Bioinformatics 2017, 33, (21), 3471–3472.

87. MacLean, B.; Tomazela, D. M.; Shulman, N.; Chambers, M.; Finney, G. L.; Frewen, B.; Kern, R.; Tabb, D. L.; Liebler, D. C.; MacCoss, M. J., Skyline: an open source document editor for creating and analyzing targeted proteomics experiments. Bioinformatics 2010, 26, (7), 966–8.

88. Reiter, L.; Rinner, O.; Picotti, P.; Huttenhain, R.; Beck, M.; Brusniak, M. Y.; Hengartner, M. O.; Aebersold, R., mProphet: automated data processing and statistical validation for large-scale SRM experiments. Nat Methods 2011, 8, (5), 430–5.

89. Govaert, E.; Van Steendam, K.; Willems, S.; Vossaert, L.; Dhaenens, M.; Deforce, D., Comparison of fractionation proteomics for local SWATH library building. Proteomics 2017, 17, (15–16).

90. Navarro, P.; Kuharev, J.; Gillet, L. C.; Bernhardt, O. M.; MacLean, B.; Röst, H. L.; Tate, S. A.; Tsou, C. C.; Reiter, L.; Distler, U.; Rosenberger, G.; Perez-Riverol, Y.; Nesvizhskii, A. I.; Aebersold, R.; Tenzer, S., A multicenter study benchmarks software tools for label-free proteome quantification. Nat Biotechnol 2016, 34, (11), 1130–1136.

91. Tully, B.; Balleine, R. L.; Hains, P. G.; Zhong, Q.; Reddel, R. R.; Robinson, P. J., Addressing the Challenges of High-Throughput Cancer Tissue Proteomics for Clinical Application: ProCan. Proteomics 2019, 19, (21-22), e1900109.

92. Aggarwal, S.; Yadav, A. K., False Discovery Rate Estimation in Proteomics. Methods Mol Biol 2016, 1362, 119–28.

93. Buscail, E.; Alix-Panabières, C.; Quincy, P.; Cauvin, T.; Chauvet, A.; Degrandi, O.; Caumont, C.; Verdon, S.; Lamrissi, I.; Moranvillier, I., High Clinical Value of Liquid Biopsy to Detect Circulating Tumor Cells and Tumor Exosomes in Pancreatic Ductal Adenocarcinoma Patients Eligible for Up-Front Surgery. Cancers 2019, 11, (11), 1656.

94. Voskuil, J., Commercial antibodies and their validation. F1000Research 2014, 3, 232–232.

95. Anderson, N. L.; Anderson, N. G., The human plasma proteome: history, character, and diagnostic prospects. Mol Cell Proteomics 2002, 1, (11), 845–67.

96. Doerr, A., DIA mass spectrometry. Nature Methods 2015, 12, (1), 35–35.

97. Zhang, F.; Ge, W.; Ruan, G.; Cai, X.; Guo, T., Data-Independent Acquisition Mass Spectrometry-Based Proteomics and Software Tools: A Glimpse in 2020. Proteomics 2020, e1900276.

98. Doellinger, J.; Blumenscheit, C.; Schneider, A.; Lasch, P., Isolation Window Optimization of Data-Independent Acquisition Using Predicted Libraries for Deep and Accurate Proteome Profiling. Anal Chem 2020.

99. Shen, J.; Pagala, V. R.; Breuer, A. M.; Peng, J.; Bin, M.; Wang, X., Spectral Library Search Improves Assignment of TMT Labeled MS/MS Spectra. J Proteome Res 2018, 17, (9), 3325–3331.

100. Rice, S. J.; Liu, X.; Zhang, J.; Belani, C. P., Absolute Quantification of All Identified Plasma Proteins from SWATH Data for Biomarker Discovery. Proteomics 2019, 19, (3), e1800135.

101. Rosenberger, G.; Liu, Y.; Röst, H. L.; Ludwig, C.; Buil, A.; Bensimon, A.; Soste, M.; Spector, T. D.; Dermitzakis, E. T.; Collins, B. C.; Malmström, L.; Aebersold, R., Inference and quantification of peptidoforms in large sample cohorts by SWATH-MS. Nat Biotechnol 2017, 35, (8), 781–788.

102. Collins, B. C.; Hunter, C. L.; Liu, Y.; Schilling, B.; Rosenberger, G.; Bader, S. L.; Chan, D. W.; Gibson, B. W.; Gingras, A. C.; Held, J. M.; Hirayama-Kurogi, M.; Hou, G.; Krisp, C.; Larsen, B.; Lin, L.; Liu, S.; Molloy, M. P.; Moritz, R. L.; Ohtsuki, S.; Schlapbach, R.; Selevsek, N.; Thomas, S. N.; Tzeng, S. C.; Zhang, H.; Aebersold, R., Multi-laboratory assessment of reproducibility, qualitative and quantitative performance of SWATH-mass spectrometry. Nat Commun 2017, 8, (1), 291.

103. Ahn, S. B.; Mohamedali, A.; Anand, S.; Cheruku, H. R.; Birch, D.; Sowmya, G.; Cantor, D.; Ranganathan, S.; Inglis, D. W.; Frank, R.; Agrez, M.; Nice, E. C.; Baker, M. S., Characterization of the interaction between heterodimeric αvβ6 integrin and urokinase plasminogen activator receptor (uPAR) using functional proteomics. J Proteome Res 2014, 13, (12), 5956–64.

104. Guo, H.; Ling, C.; Ma, Y. Y.; Zhou, L. X.; Zhao, L., Prognostic role of urokinase plasminogen activator receptor in gastric and colorectal cancer: A systematic review and meta-analysis. Onco Targets Ther 2015, 8, 1503–9.

105. Liu, K. L.; Fan, J. H.; Wu, J., Prognostic Role of Circulating Soluble uPAR in Various Cancers: a Systematic Review and Meta-Analysis. Clin Lab 2017, 63, (5), 871–880.

106. Pizzatti, L.; Panis, C.; Lemos, G.; Rocha, M.; Cecchini, R.; Souza, G. H.; Abdelhay, E., Label-free MSE proteomic analysis of chronic myeloid leukemia bone marrow plasma: disclosing new insights from therapy resistance. Proteomics 2012, 12, (17), 2618–31.

