## Supplementary material for "A Recombinant Protein Biomarker DDA Library Increases DIA Coverage of Low Abundance Plasma Proteins": Skyline peptide and transition settings.

**Supplementary Table S1**: Skyline peptide and transition settings.

| **Integrate All** | | |
| --- | --- | --- |
| ***Peptide Settings*** | | |
| Digestion | Enzyme | Trypsin [KR \| P] |
|  | Max missed cleavages | 0 |
|  | Background proteome | None |
| Prediction | Retention time predictor | Created |
|  | Use measured retention time when present | checked |
|  | Time window(min) | 2 |
|  | Ion mobility predictor | None |
|  | Use spectral library drift times when present | unchecked |
|  | Resolving power | (blank) |
| Filter | Min length | 7 |
|  | Max length | 36 |
|  | Exclude N terminal Aas | 36 |
|  | Exclude potential ragged ends | unchecked |
|  | Exclude peptides containing | (blank) |
|  | Auto-select all matching peptides | checked |
| Library | Library | Created |
|  | Pick peptides matching | Library |
|  | Rank peptides by | Picked Intensity |
|  | Limit peptides per proteins | (blank) |
|  | Peptides | (blank) |
| Modifications | Structural modifications | Carbamidomethyl (C), Oxidation (M), Water Loss (D, E, S, T) |
|  | Max variable mods | 3 |
|  | Max neutral losses | 1 |
|  | Isotope label type | heavy |
|  | Isotope modifications | None |
|  | Internal standard type | heavy |
| Quantification | Normalization Method | Equalize Medians |
|  | All others | None |
| ***Transition Settings*** | |  |
| Prediction | Precursor mass | Monoisotopic |
|  | Product ion mass | Monoisotopic |
|  | Collision Energy | None |
|  | Declustering potential | None |
|  | Optimization library | None |
|  | Compensation voltage | None |
|  | Use optimization values when present | unchecked |
| Filter | Precursor charges | 2, 3 |
|  | Ion charges | 1, 2, 3 |
|  | Ion types | y, b |
|  | Product ions from | ion 3 |
|  | Product ions to | last ion |
|  | Product ions - Special ions | N-terminal to Proline |
|  | Use DIA precursor window for exclusion | checked |
|  | Auto select all matching transitions | checked |
| Library | Ion match tolerance | 0.5 |
|  | If a library spectrum is available, pick its most intense ions | checked |
|  | Pick XX product ions | 6 |
|  | Pick XX minimum product ions | 6 |
|  | From filtered ions charges and types | unchecked |
|  | From filtered ions charges and types plus filtered product ions | unchecked |
|  | From filtered product ions | checked |
| Instrument | Min m/z | 350 |
|  | Max m/z | 2000 |
|  | Dynamic min product m/z | unchecked |
|  | Method match tolerance m/z | 0.055 |
|  | Firmware | (blank) |
|  | Firmware | (blank) |
|  | Min time | (blank) |
|  | Max time | (blank) |
| Full-Scan | MS1 - Isotope peaks included | None |
|  | MS1 - Precursor mass analyzer | (blank) |
|  | MS1 - Peaks | (blank) |
|  | MS1- Resolving power | None |
|  | MS1- Isotope labeling enrichment | (blank) |
|  | MS/MS- Acquisition method | DIA |
|  | MS/MS- Product mass analyzer | TOF |
|  | MS/MS- Isolation scheme | SWATH wiff file |
|  | MS/MS Resolving power | 30,000 |
|  | Retention time filtering - Use only scans within XX min of MS/MS IDs | unchecked |
|  | Retention time filtering - Use only scans within XX min of predicted RTs | 5 |
|  | Retention time filtering - Include all matching scans | unchecked |
| ***Document Settings*** | | |
| Annotations | | None |
| Group Comparisons | | None |
